# Global view of domain-specific O-linked mannose glycosylation in glycoengineered cells

**DOI:** 10.1101/2024.01.15.575371

**Authors:** Lorenzo Povolo, Weihua Tian, Sergey Y. Vakhrushev, Adnan Halim

**Affiliations:** Copenhagen Center for Glycomics, Department of Cellular and Molecular Medicine, Faculty of Health and Medical Sciences, University of Copenhagen, Blegdamsvej 3B, DK-2200 Copenhagen N, Denmark; Department of Biotechnology and Biomedicine, Technical University of Denmark, Søltofts Plads, Building 224, DK-2800 Kgs. Lyngby, Denmark

**Keywords:** Glycobiology, post-translational modification (PTM), O-Man, Protein O-mannosyltransferases, mass spectrometry (MS), lectins, genetic engineering, CRISPR/Cas, protein domains, cadherin, plexin, dystroglycan

## Abstract

Protein O-linked mannose (O-Man) glycosylation is an evolutionary conserved post-translational modification (PTM) that fulfills important biological roles during embryonic development. Three non-redundant enzyme families, POMT1/POMT2, TMTC1-4 and TMEM260, selectively coordinate the initiation of protein O-Man glycosylation on distinct classes of transmembrane proteins, including α-dystroglycan, cadherins and plexin receptors. However, a systematic investigation of their substrate specificities is lacking, in part due to the ubiquitous expression of O-Man glycosyltransferases in cells, which precludes analysis of pathway-specific O-Man glycosylation on a proteome-wide scale. Here, we apply a targeted workflow for membrane glycoproteomics across five human cell lines to extensively map O-Man substrates and genetically deconstruct O-Man initiation by individual and combinatorial knock-out (KO) of O-Man glycosyltransferase genes. We established a human cell library for analysis of substrate specificities of individual O-Man initiation pathways by quantitative glycoproteomics. Our results identify 180 O-Man glycoproteins, demonstrate new protein targets for the POMT1/POMT2 pathway and show that TMTC1-4 and TMEM260 pathways widely target distinct Ig-like protein domains of plasma membrane proteins involved in cell-cell and cell-extracellular matrix interactions. The identification of O-Man on Ig-like folds adds further knowledge on the emerging concept of domain-specific O-Man glycosylation which opens for functional studies of O-Man glycosylated adhesion molecules and receptors.

## Introduction

Protein O-linked mannose glycosylation (O-Man) at serine (Ser) and threonine (Thr) residues is an essential protein post-translational modification (PTM) conserved from bacteria to humans (1–4). Biosynthesis of O-Man is initiated in the endoplasmic reticulum (ER) lumen by integral transmembrane GT-C_A_ enzymes (5–7) that utilize lipid-linked dolichol phosphate mannose (Dol-P-Man) as donor substrate (8). In yeast, seven protein O-mannosyltransferases (PMT1-PMT7) are known to orchestrate O-Man initiation on at least 25% of the proteins that traffic the ER and secretory pathway (9,10). The mammalian orthologs POMT1 and POMT2, which are absent in plants, are distinguished from yeast PMTs by their narrow substrate specificities and dedicated functions for O-Man initiation on only a few human proteins, including KIAA1549, SUCO and α-dystroglycan (α-DG) (11–13). Mammals have evolved a complex biosynthetic machinery involving at least 17 genes, among which *POMT1, POMT2, POMGNT1, POMGNT2, MGAT5B, B3GALNT2, FKTN, FKRP, TMEM5 (RXYLT1), B4GAT1, LARGE1* and *LARGE2* participate in the assembly of complex O-Man glycans on α-DG of the dystrophin-associated glycoprotein complex (DGC) (14–16). Functional O-Man glycosylation of α-DG is required for interactions with extracellular matrix (ECM) components, including laminin, agrin and perlecan, which are anchored to the DGC and the actin cytoskeleton through a complex O-Man polysaccharide known as matriglycan (14).

More recently, we have shown that mammals have evolved multiple biosynthetic pathways for O-Man initiation based on the GT-C_A_ type enzymes TMTC1, TMTC2, TMTC3, TMTC4 and TMEM260 (17–20). The TMTC1-4 and TMEM260 enzymes, classified in the CAZy database as GT105 and GT117, respectively, serve distinct classes of extracellular Ig-like protein domains found on transmembrane adhesion molecules (cadherins) and receptors (plexins) (17,18). Unlike α-DG, O-Man on Ig-like domains is not elongated into complex structures but is rather present as single *α*-linked mannose monosaccharides on β-strands of the Ig-like domains. The TMTC1-4 enzymes are dedicated to the cadherin superfamily of adhesion molecules and selectively glycosylate highly conserved Ser/Thr residues with O-Man in two β-strands (B and G) of extracellular cadherin (EC) domains (17), which are functional units that direct homo/heterophilic *cis-* and *trans*-binding important for cell-cell interactions (21,22). The recently identified TMEM260 enzyme selectively serves extracellular immunoglobulin, plexin, transcription factor (IPT) domains found among a subset of structurally related plasma membrane receptors, including hepatocyte growth factor receptor (cMET), macrophage-stimulating protein receptor (MST1R or Recepteur d’Origine Nantais, RON) and members of the plexin family (17,18). The IPT domain is a distinct subclass of the Ig-like fold, and while the functions of their O-Man glycans, located on conserved Ser/Thr residues within β-strands of IPT domains, remains unknown, TMEM260-driven O-Man glycosylation appears to be critical for receptor maturation and epithelial morphogenesis (18).

O-Man glycosylation fulfils important roles during development, and dysregulation of O-Man initiation is associated with severe developmental disorders in humans. Mutations in *POMT1/POMT2* genes is linked to a subclass of congenital muscular dystrophies known as α-dystroglycanopathies, characterized by progressive muscular degeneration and developmental abnormalities in brain and eyes (23–25). Genetic defects in *TMTC1-4* are primarily associated with neurological disorders, including brain malformation and hearing loss (26–30) while bi-allelic mutations in *TMEM260* underlie the SHDRA syndrome, characterized by congenital heart defects, kidney phenotypes and neurodevelopmental disorders (18,31,32).

However, the specific contributions of individual biosynthetic enzymes and O-Man initiation pathways towards the modified glycoproteome, as well as the mechanisms underlying substrate selection and potential crosstalk, remain understudied. The lack of understanding is in part due to the widespread and ubiquitous expression of O-Man glycosyltransferases across cell types (33), which precludes studies of individual biosynthetic pathways without interference from isoenzymes or other O-Man glycosyltransferases. A comprehensive investigation of enzyme specificities and their substrate selectivity may thus not only improve the understanding of receptor functions and regulations, but also uncover molecular details that explain phenotypes arising from pathway-specific dysfunctions in O-Man glycosylation. In this study, we therefore aimed to dissect the biosynthetic regulation and pathway-specific O-Man initiation for POMT1/POMT2, TMTC1-4 and TMEM260 enzyme families.

We established an improved O-glycoproteomics workflow for targeted analysis of transmembrane O-Man substrates in five human cancer cell lines representing different tissue origins and expanded the human O-Man glycoproteome. Furthermore, we established a panel of genetically engineered human HEK293 cell lines with defined O-Man glycosylation capacities for differential O-Man glycoproteomics. We identify 180 O-Man glycoproteins (of which 67 not previously described) and dissect individual contributions of POMT1/POMT2, TMTC1-4 and TMEM260 enzymes. We report the hitherto largest O-Man glycoproteome with site-specific mapping across a wide range of protein classes involved in ECM/cell-cell interactions, which opens for further functional studies of distinct protein classes based on their O-Man glycosylation status.

## Results

### An optimized workflow for O-Man glycoproteomics

Sensitive glycoproteomics largely hinges on optimized preparation, efficient glycopeptide enrichment and precise quantification of the analyzed sample(s) (34). We first aimed to improve these three key steps in our established workflow (13,17) for identification and quantification of O-Man proteins and establish an optimized workflow for O-Man glycoproteomics. Previous glycoproteomics studies have demonstrated that the majority (72%) of proteins undergoing O-Man glycosylation consists of single-pass or multi-pass transmembrane proteins (13,17–19,35) (**Fig. S1A**). To improve sensitivity in O-Man glycoproteomics analysis, we therefore envisioned that crude membrane preparation (CMP) (36) would enrich the transmembrane sub-proteome and exclude abundant cytosolic proteins that are typically extracted in total cell lysates (TCL). We separately analyzed HEK293 TCL and CMP samples by detergent-assisted protein extraction, proteolytic digestion and lectin enrichment of O-Man glycopeptides (**Fig. 1A**). Bottom-up analyses of HEK293 cells subjected to CMP extraction identified 520 O-Man glyco-PSMs originating from 54 O-Man glycoproteins, while only 275 O-Man glyco-PSMs from 39 unique O-Man glycoproteins were detected in the same sample subjected to TCL extraction (**Fig. 1B**), thus demonstrating that CMP extraction improves analytical sensitivity for O-Man glycoproteomics. Second, we sought to improve O-Man glycopeptide enrichment from complex mixtures by using the α-mannose-specific *Burkholderia cenocepacia* lectin A (BC2L-A) (37,38). BC2L-A, a 28 kDa dimeric and Ca^2+^-dependent lectin capable of binding both O-Man and C-linked mannose glycans on tryptophan residues (C-Man), has emerged as an alternative to ConA for improved enrichment of glycopeptides by LWAC (35). Moreover, BC2L-A was found to capture O-Man glycopeptides in ConA-LWAC flow-through fractions, clearly indicating that BC2L-A is a better affinity reagent for O-Man glycoproteomics compared to ConA (35). Therefore, we chose to substitute ConA with the BC2L-A lectin in our LWAC procedure for improved O-Man glycopeptide enrichment.

**Figure 1.**
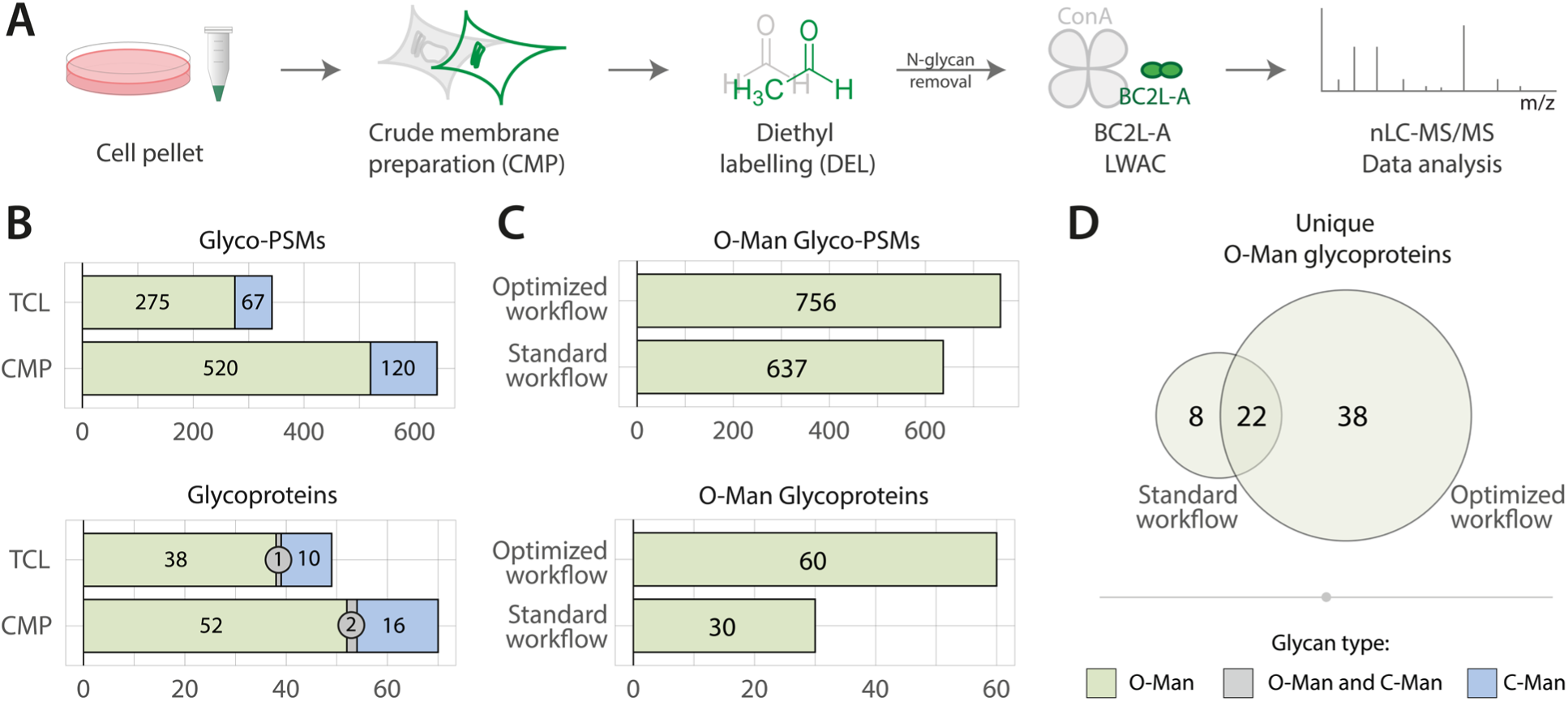
An optimized workflow for O-Man glycoproteomics. **(A)** Schematics of the optimized workflow, including crude membrane preparation of cells (CMP), diethyl labelling (DEL), N-glycan removal, BC2L-A LWAC and nLC-MS/MS. Key steps for the optimized workflow are highlighted in green. **(B)** Comparison of glycopeptide enrichment of HEK293 total cell lysate (TCL) and crude membrane preparations (CMP). CMP leads to an increase of both glyco-PSMs and glycoproteins detected in this test run experiment. Color code as in *panel (D)*. **(C)** Comparison of O-Man data identified in one experiment run with *standard workflow* in our previous work (17) and the ones with the *improved workflow* employed in this work. For this information, only dimethyl medium or diethyl heavy channels of the same cell line were considered. The current workflow allows a 2x identification of unique O-Man proteins. **(D)** Comparison of O-Man data of panel (C) in terms of unique proteins identified. More than 4x unique proteins were identified exclusively with the optimized compared to the standard workflow.

Furthermore, we assessed the approach for differential labelling of peptides by stable isotopes. In our most recent study (18), we employed diethyl labeling (DEL) for two-plex experiments, which is a cost-effective, flexible, and readily available method that utilizes ^13^C isotopologues of acetaldehyde, as an alternative to dimethyl labelling (DML) (39). In agreement with previous studies (39), the use of diethyl labelling improved quantification accuracy and retention of short hydrophilic O-Man glycopeptides on C18 columns (**Fig. S1B**). Diethyl labelling of peptides has also been reported to enhance ionization of hydrophilic peptides (39), thus improving the number of identifications in bottom-up analyses. We therefore also employed light DEL in one-plex experiments where no quantification was necessary, to improve the sensitivity in our glycoproteomic workflows.

Taken together, the optimized workflow combining CMP, DEL and BC2L-A enrichment (**Fig. 1A**) enabled significant improvements in bottom-up O-Man glycoproteomics, with a 2-fold increase in total number of O-Man proteins identified in comparison to the standard method used in our previous studies (17) (**Fig. 1C**). The optimized workflow (**Fig 1D**) allowed identification of 60 O-Man glycoproteins in total (n=38 exclusive) compared to the standard workflow (17), which identified 30 O-Man glycoproteins (n=8 exclusive), clearly demonstrating the overall improved sensitivity for O-Man glycoproteomics.

### An expanded O-Man glycoproteome in five human cells lines

We analyzed five human cancer cell lines derived from different tissues, including BG1 (ovarian adenocarcinoma), CaCo-2 (colorectal adenocarcinoma), HEK293 (embryonic kidney), HepG2 (hepatocellular carcinoma) and SH-SY5Y (neuroblastoma), using the optimized glycoproteomics workflow described above (**Fig. 1A**). CMP tryptic digests from each cell line were labelled with light DEL before O-Man glycopeptide enrichment by BC2L-A LWAC and mass spectrometry. HCD/ETciD-based bottom-up analyses identified 3882 glyco-PSMs from 120 unique O-Man proteins (**Fig. 2A**), 33 of which have not been described as O-Man substrates before (13,17–19,35). Notably, we identified 79 O-Man glycoproteins in HEK293 cells alone, which represents a two-fold improvement compared to our standard workflow (17), further demonstrating the improved sensitivity of the method used here. Gene ontology enrichment analysis of the 120 proteins identified showed strong enrichment of biological process terms for cell adhesion (**Fig. 2B**), in agreement with the function of canonical substrates for O-Man proteins such as cadherins and protocadherins (20). Indeed, cadherins and protocadherins constitute the largest class of O-Man proteins identified in these datasets, where single O-Man monosaccharides modify extracellular cadherin (EC) domains of 54 unique members of the cadherin superfamily. While some O-Man proteins appear to be cell-line specific, the majority of the O-Man proteins were detected in at least two different cell lines (**Fig. 2C**). As recently described, IPT domains are TMEM260 substrates (18), and here, we identified 7 members (PLXNA1-4, PLXNB2-3, PLXND1) of the plexin family, as well as cMET and RON receptors, which account to 9 proteins (of n=12 predicted) with O-Man on IPT domains. Interestingly, we also identified 26 proteins with O-Man on distinct Ig-like domains, including Ig-like C2-type domains (n=5), Fibronectin type-III domains (n=4) and Ig-like V-type domains (n=2), indicating that O-Man glycosyltransferases selectivity is not limited to the Ig-like fold subclasses defined by EC- and IPT-domains. While most of the identified O-Man glycopeptides map to annotated protein domains, 34 additional proteins were identified with O-Man glycosylation in unstructured/unannotated regions as reported in Uniprot. As expected, O-Man glycosylation in unstructured protein domains was identified on α-DG, KIAA1549 and SUCO, which are substrates of the classical POMT1/POMT2 pathway (13), but also on a subset of proteins that have not been previously reported as O-Man targets, mainly representing ER-localized or membrane proteins. In the five human cell lines examined, we identified the largest and most diverse set of O-Man proteins in HEK293 cells. We therefore chose to glycoengineer HEK293 cells by CRISPR/Cas9 KO to deconstruct O-Man initiation pathways and establish suitable cell models for differential O-Man glycoproteomics.

**Figure 2.**
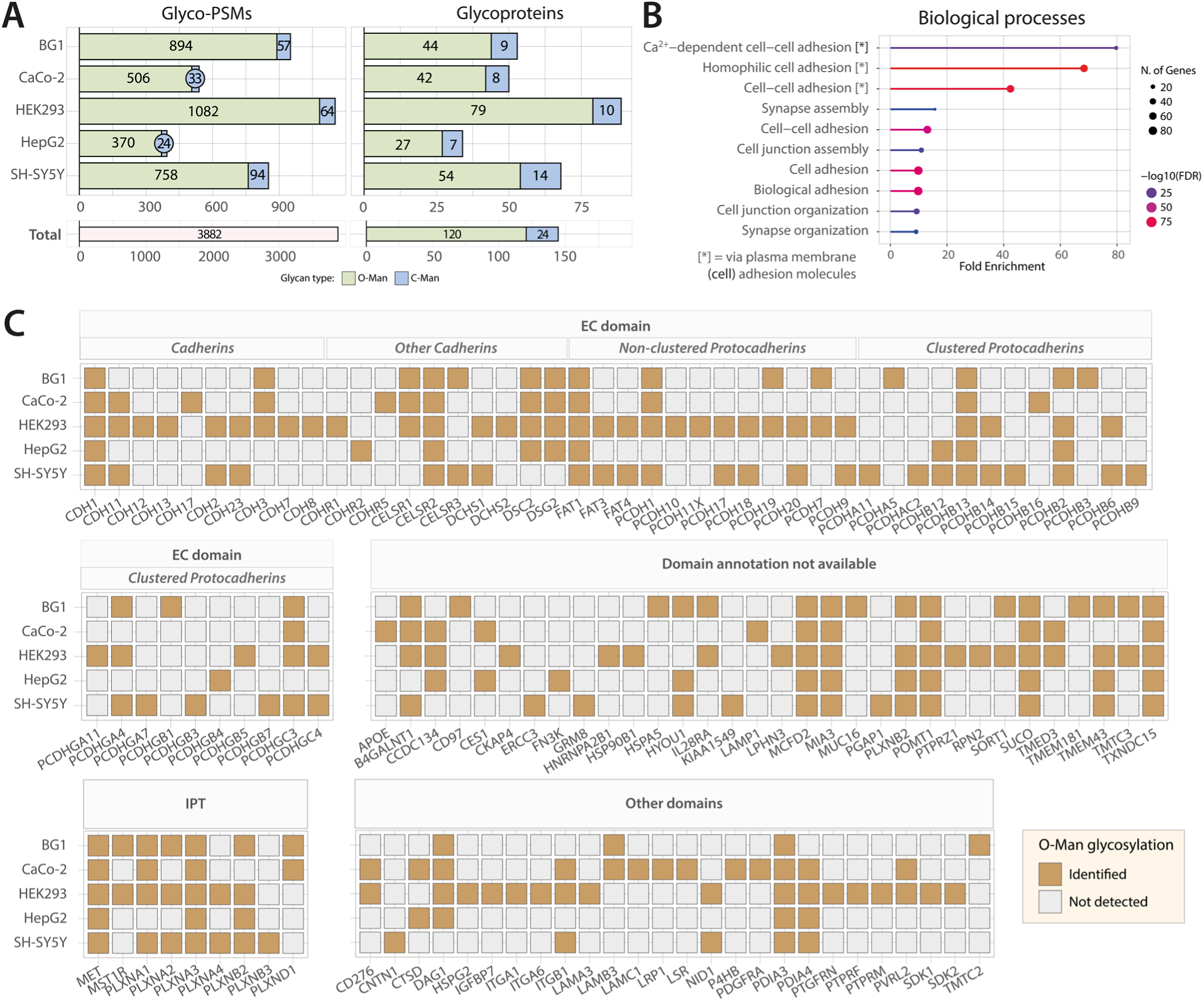
The O-Man glycoproteome of five mammalian cell lines. **(A)** Comparison of glyco-PSMs and glycoproteins identified across the five different cell lines analyzed. HEK293 data are more abundant in O-Man glyco-PSMs and glycoproteins, while data from SH-SY5Y cells contains more C-Man proteins. **(B)** Gene ontology enrichment analysis for biological processes underlying the proteins identified in the datasets. Cell adhesion appears to be the most enriched term across the O-Man hits identified. **(C)** Overview of identified proteins across cell lines. Proteins are divided into groups based on domains (EC, IPT, Other domains, Domain annotation not available) according to the Uniprot database (May 2022). Proteins containing EC domains are subdivided according to cadherins/protocadherin classes. Ocher color indicates that the protein has been identified with at least one O-Man glycosylated residue on the respective annotated domain. Gray signifies that the protein in the respective domain has not been detected with O-Man sugars in the specific cell line.

### O-Man glycosyltransferase specificities in genetically deconstructed HEK293 cells

To study specificities, relationships, and potential crosstalk between the POMT1/POMT2, TMTC1-4 and TMEM260 glycosyltransferase families responsible for initiation of O-Man biosynthesis, we generated a library of HEK293 cell lines with combinatorial CRISPR-Cas9 knock-out of selected genes for a partial and complete deconstruction of O-Man glycosylation capacity in HEK293 cells (**Fig. 3A**). The genetic engineering was undertaken in the HEK293 *COSMC/POMGNT1* KO (SimpleCell; HEK293^SC^) background (13,40), which are not capable of producing core-M1 or -M2 type complex O-Man glycans (41,42), thus allowing a simplified one-step (BC2L-A LWAC) enrichment and identification of O-Man glycosites on diverse protein classes. To investigate pathway-specific O-Man initiation, we employed a combinatorial KO design in HEK293^SC^ cells to inactivate two out of three enzyme families in combination, which allowed us to establish individual cell lines where either *POMT1/POMT2, TMTC1-4* or *TMEM260* genes remained unedited. For simplicity, we hereafter refer to each HEK293 cell line by the active biosynthetic pathway: *e.g.,* a combinatorial KO of *COSMC/POMGNT1/POMT1/POMT2/TMTC1-4* genes in HEK293 is denoted as “HEK293^TMEM260^” (**Fig. 3A**). For control experiments, we also established a HEK293 cell line with complete deconstruction of O-Man glycosylation capacity by *COSMC/POMGNT1/POMT1/POMT2/TMTC1-4/TMEM260* KO, which we here refer to as the “HEK293^nO-Man^” cell line.

**Figure 3.**
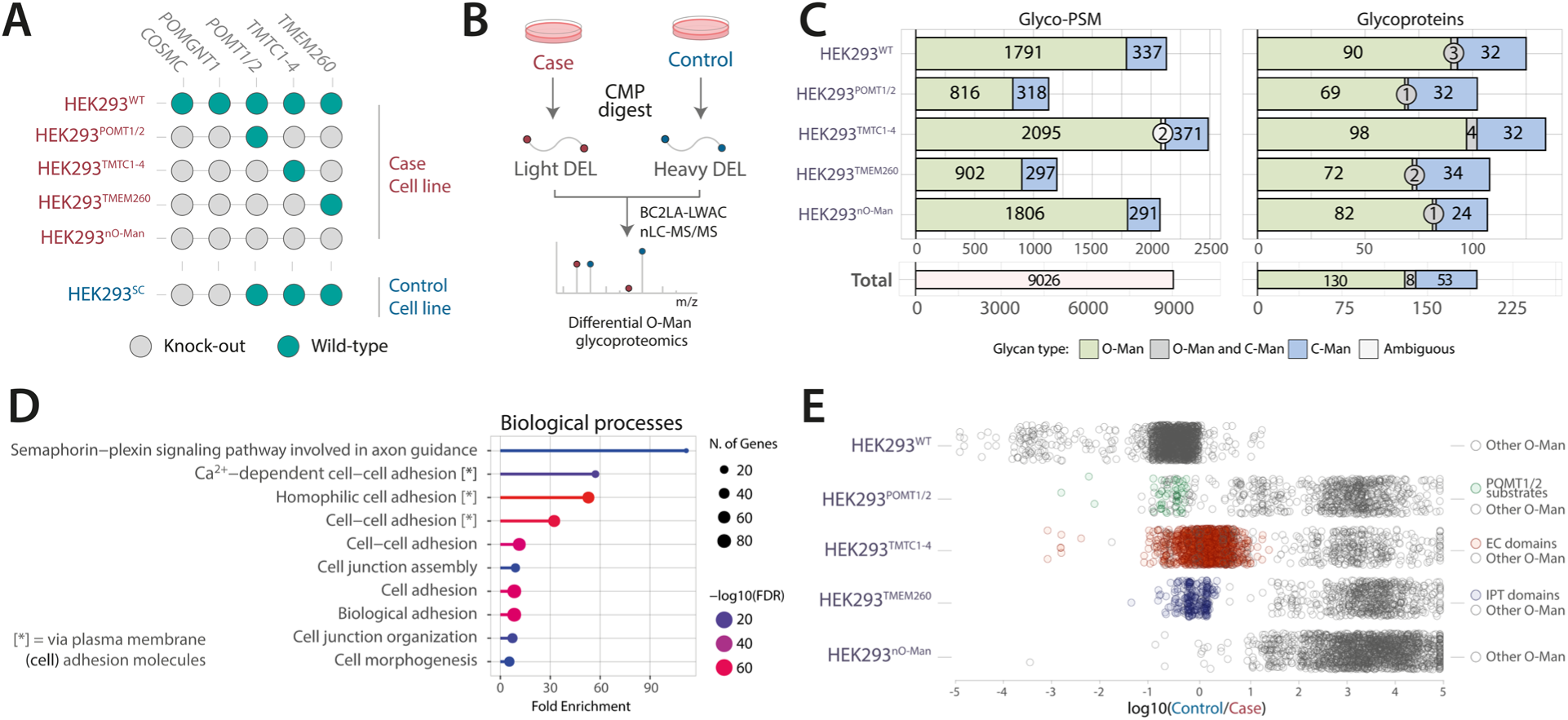
MS analysis of a glycoengineered HEK293 cell panel. **(A)** Graphical depiction of glycoengineered cell matrix with cell line identifier on the left, and genes targeted by genetic engineering on top (italic). **(B)** Schematics for differential O-Man glycoproteomics used in this study, based on the optimized workflow in Figure 1A. **(C)** Glyco-PSMs and glycoproteins identified across these five datasets. **(D)** Gene ontology enrichment analysis for biological processes in which O-Man proteins identified are involved; terms regarding cell-cell adhesion show the highest enrichment. **(E)** Glycoengineered cells where only one pathway is left unaltered show O-Man glycosylation capacity like the HEK293^SC^ control, in line with the previous knowledge on canonical O-Man substrates. Each circle represents a O-Man PSM, and log_10_(Heavy/Light)=0 signifies that the O-Man capacity for the specific PSM in glycoengineered cell is equal to WT. Only the canonical substrates are colored in each glycoengineered cell line for clarity. **(F)** HEK293^WT^ cells show O-Man glycosylation capacity like HEK293^SC^, confirming that the HEK293^SC^ is a suitable control cell line for this experiment. The HEK293^nO-Man^ cell line shows global loss of O-Man glycosylation compared to HEK293^SC^ control. Each circle represents a O-Man PSM, and log_10_(Heavy/Light)=0 signifies that the O-Man capacity for the specific PSM in glycoengineered cell is equal to the one of HEK293^SC^.

To study enzymes specificities, we applied the optimized workflow for differential glycoproteomics analyses (**Fig. 3B**) of HEK293^TMEM260^, HEK293^TMTC1-4^, HEK293^POMT1/POMT2^ and HEK293^nO-Man^ for comparative analysis using HEK293^SC^ as control. We also performed a differential analysis comparing HEK293^WT^ and HEK293^SC^ to assess if *COSMC/POMGNT1* KO influenced O-Man initiation (**Fig. 3A**). Collectively, the five comparative datasets identified a total of 9026 glyco-PSMs, 7410 of which were O-Man and 1614 were C-Man PSMs (**Fig. 3C**). A Gene Ontology Enrichment Analysis was performed on the 138 unique O-Man proteins identified, suggesting that biological process terms related to cell signaling and adhesion were enriched, in agreement with the functions of cadherins and plexins. (**Fig. 3D**). Proteomics of heavy and light labelled tryptic digests analyzed after 1:1 mixing revealed a normal distribution of H/L ratios for case/control samples, and we used interquartile range (IQR) calculations for each comparison to determine biological/technical variability and outliers by Q1-1.5xIQR and Q3+1.5xIQR boundaries (**Fig. S2A**), which formed the basis for setting a ±10-fold change as a threshold for differential regulation of O-Man glycosylation in case/control comparisons.

We proceeded with analyses of differentially labelled O-Man glycopeptides from case/control comparisons to evaluate O-Man glycoproteome changes in the engineered cell lines (**Fig. 3E**), and first examined relative abundances of O-Man glycopeptides in the differential analysis of HEK293^WT^ / HEK293^SC^ cells. We observed that >90% of the O-Man glycopeptide H/L ratios were within the technical variability (±10-fold) of the measurement and could thus conclude that there is no significant difference in O-Man initiation between HEK293^WT^ and HEK293^SC^ cells (**Fig. 3E**). For HEK293^POMT1/POMT2^, HEK293^TMTC1-4^ and HEK293^TMEM260^, which were all compared to O-Man initiation in the HEK293^SC^ background, we observed H/L ratio spread across 6 orders of magnitude in all comparisons, indicating that combinatorial KOs in each cell line influenced O-Man initiation of distinct substrates. More specifically, in the HEK293^POMT1/POMT2^ / HEK293^SC^ comparison, we observed that O-Man glycopeptides from KIAA1549, SUCO and α-DG did not change in relative abundance (±10-fold), thus demonstrating that POMT1/POMT2 is solely responsible for initiation of O-Man biosynthesis within unstructured regions of KIAA1549, SUCO and α-DG. In contrast, O-Man glycopeptides derived from cadherins or plexins where >100 fold more abundant in the HEK293^SC^ cell line (**Supplementary material**), which demonstrated that HEK293^POMT1/POMT2^ cells are unable to induce O-Man initiation on EC-domains (TMTC1-4 dependent) or IPT-domains (TMEM260 dependent). Furthermore, we confirmed TMTC1-4 dependent O-Man initiation of EC-domains in the HEK293^TMTC1-4^ / HEK293^SC^ differential analysis, where O-Man glycopeptides derived from cadherins (n=1453) were found to be equally abundant while selective loss of O-Man was observed on KIAA1549, SUCO and α-DG (POMT1/POMT2 dependent) and on IPT-domains (**Fig. 3E**, **Supplementary material**). Moreover, the differential analysis of HEK293^TMEM260^ / HEK293^SC^ cells demonstrated loss of O-Man glycosylation on cadherins, KIAA1549, SUCO and α-DG while no changes (±10-fold) were observed for O-Man glycopeptides derived from IPT-domains (**Fig. 3E**). Finally, in the HEK293^nO-Man^ / HEK293^SC^ comparison, we observed a screwed distribution of H/L ratios for O-Man glycopeptides, with >98% of the ratios found above the 10-fold change threshold, thus demonstrating that O-Man glycosylation is abolished upon *POMT1/POMT2/TMTC1-4/TMEM260* KO (**Fig. 3E**). Taken together, these results, which agree with previous studies (13,17,18), validated the approach and showed that our O-Man deconstruction cell library is suitable for analyses of pathway-specific substrate specificities.

### The O-Man glycoproteome in the context of three biosynthetic pathways

The use of glycoengineered cells and targeted glycoproteomics workflow allowed us to map the most extensive human O-Man glycoproteome (**Supplementary material, Fig. 2, Figure S5**). We sought to compile the glycoproteomics results presenting all detected O-Man substrates in the context of their biosynthetic pathways and grouped by protein domains (**Fig. 4**), which includes the datasets from HEK293^SC^ / HEK293^nO-Man^ originally collected for the comparison between CMP and TCL, as presented in **Fig. 1B**. The differential glycoproteomics workflow allows pairwise assessment with the HEK293^SC^ control and results can be grouped into four distinct outcomes for each domain type within every detected protein. 1) “Glycosylated” describes a particular protein domain in which all glyco-PSMs detected exhibit glycosylation in both the HEK293^SC^ control and the case cell line at a similar ratio. This suggests that the genetically engineered cell line closely resembles a HEK293^SC^ cell line. Conversely, 2) “Not glycosylated” refers to a type of domain where O-Man glycosylation was found in the control cell line but not in the case cell line. This suggests that the glycoengineering resulted in the loss of O-Man glycosylation from all O-Man PSMs detected in the control cell line for each specific type of domain, within a given protein. We observed some cases, referred to as 3) “Ambiguous” in which protein domains within a protein displayed at least one O-Man PSM with glycosylation resembling the control cell line and at least one other displaying sensitivity to genetic engineering, or that the data output did not allow confident identification/quantification. Lastly, we classify 4) “Not detected” the protein domains in which no O-Man PSMs were detected in the dataset, but that were detected in any of the other datasets. The latter can be attributed to the inherent limitations of shotgun glycoproteomics, where specific precursor ions may not be selected for MS/MS fragmentation.

**Figure 4.**
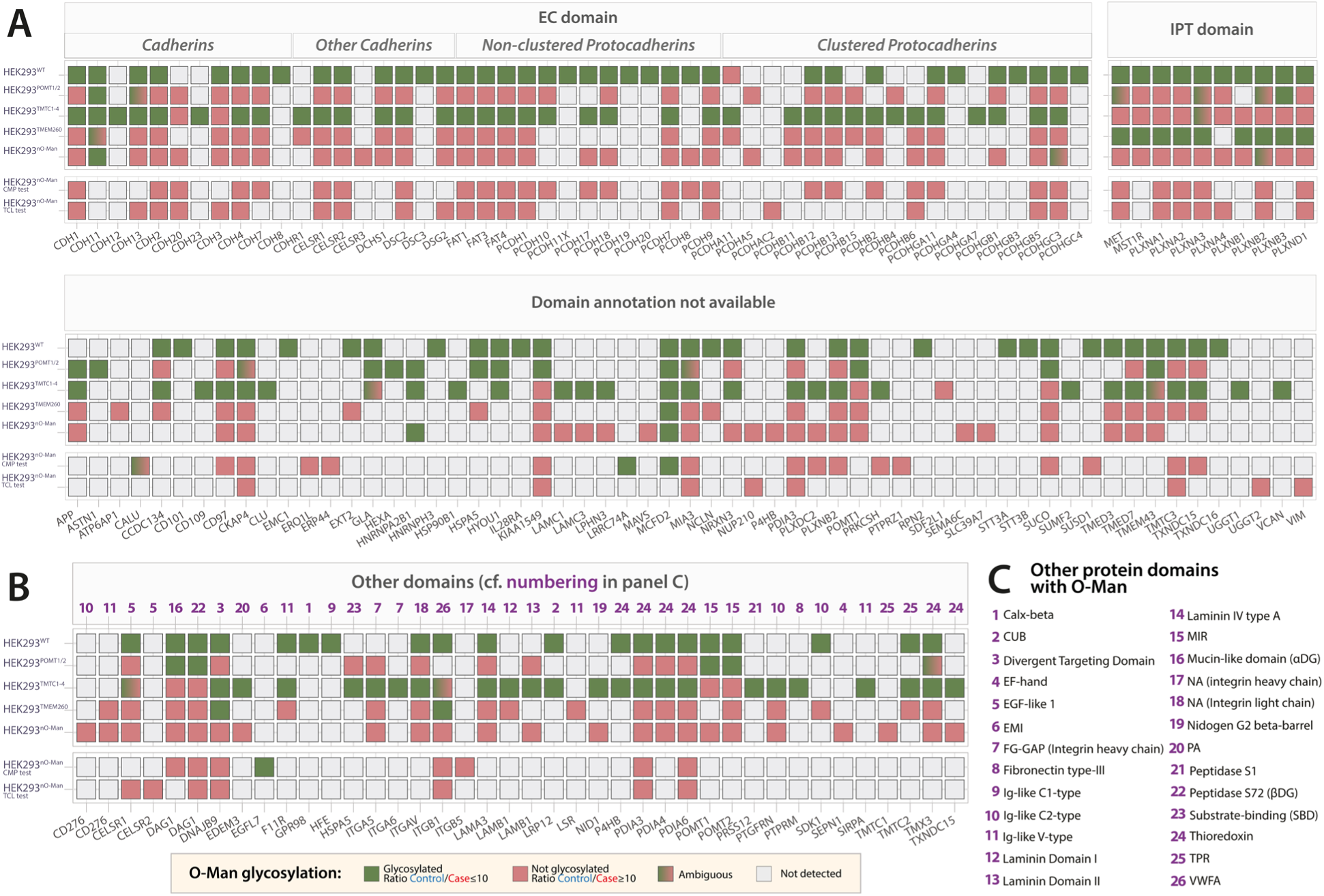
Differential O-Man glycoproteome analysis in glycoengineered HEK293 cells. **(A)** O-Man proteins identified with EC, IPT or domains without annotation. Color code is reported in *panel (B)* and clarified in detail in the Results section. **(B)** Proteins with O-Man identified on other domains, according to Uniprot annotations (May 2022). Numbering refers to the list reported in *panel (C)*. **(C)** List of domains identified with O-Man proteins. Numbering refers to proteins identified in *panel (B)*.

Among the classical O-Man substrates, we identified α-DG, SUCO, KIAA1549, as well as 50 non-classical protein substrates with O-Man on EC domains and 10 proteins with modification on IPT domains (**Fig. 4A**). HEK293^POMT1/2^ cells were, in addition to α-DG, SUCO and KIAA1549, also capable of inducing O-Man glycosylation on a limited set of other substrates, predominantly on unannotated domains or unstructured/disordered regions, including some site-specific glycosylation. Notably, HEK293^POMT1/2^ exhibit glycosylation activity within POMT1/2 MIR domains and other regions of POMT1. All proteins containing EC domains are glycosylated both in HEK293^WT^ and HEK293^TMTC1-4^ cell line, in line with previous evidence that TMTC1-4 enzymes are responsible for EC domain O-Man glycosylation (17). CDH11 and CDH13 O-Man glycosylation appears to be independent of TMTC1-4, in line with our previous study (17), and for CDH11 at least one glycosylated PSM was observed in each cell line. Additionally, while previous data (17) reported CDH13 glycosylation to be insensitive of TMTC1-4 KO, here we identify site-specific examples of glycosylation in HEK293^POMT1/2^, while HEK293^TMEM260^ cells are not capable of glycosylating EC domains of CDH13. More studies are thus warranted to elucidate the biosynthetic basis for O-Man glycosylation of the atypical CDH11 and CDH13 cadherins. Furthermore, we detect site-specific glycosylation on PCDHGC3 in HEK293^nO-Man^ dataset, but not in HEK293^POMT1/2^ nor HEK293^TMEM260^. Moreover, plexins, cMET and MST1R/RON showed O-Man glycosylation at comparable levels in HEK293^WT^ and HEK293^TMEM260^ as expected (18), as well as unreported cases of site-specific glycosylation in HEK293^POMT1/2^ (MET, PLXNA3, PLXNB2), HEK293^TMTC1-4^ (PLXNA3, PLXNB2) and in HEK293^nO-Man^ (PLXNB2). These results indicate an interplay between initiation pathways with specific proteins serving as substrates for different enzyme families, which needs further investigation. Remarkably, our platform allowed detection of O-Man glycosylation on several other proteins, both in regions with no assigned domain/unstructured regions or annotated domains (UniProt). Few of these candidates were also identified previously, such as integrins and, interestingly, a site on β-DG where O-Man glycosylation was abolished by KO of *TMEM260* (18). Here, we also identified members of α-integrins which where O-Man glycosylated in the HEK293^TMTC1-4^ cell line, while ITGB1 showed site-specific glycosylation in both HEK293^TMTC1-4^ and HEK293^TMEM260^. Overall, our results demonstrate that non-classical enzyme families (TMTC1-4 and TMEM260) target Ig-like fold and unannotated domains that show structural similarities, while POMT1/2 appear to have specificity for unstructured regions of distinct proteins.

### In-depth glycoproteomics analysis of purified reporter proteins

The identification of O-Man modifications on common protein folds prompted us to investigate the enzymatic preference and specificity in detail. We sought to use the same glycoengineered cell lines to express soluble reporter proteins and assess O-Man glycosylation and occupancy at specific protein domains. We adopted a conventional approach by recombinant expression of reporter constructs, digestion with trypsin or chymotrypsin before bottom-up mass spectrometry analyses (43). We sought to analyze representative proteins to assess O-Man glycosylation, specificity and, importantly, occupancy/stoichiometry at the site-specific level, and expressed 12 constructs encoding His-tagged reporter proteins (ectodomains) in HEK293^SC^ cells and HEK293^nO-Man^ cells. These included both canonical substrates (DAG1, CADH1, CDH11), non-canonical substrates (ITGB1, ITGA5, ITGAV, F11R), as well as proteins not identified as O-Man proteins in this study but belonging to the same family of O-Man proteins identified here (ITGA2, SEMA4D, PTPRJ) or containing protein domains identified with O-Man in this study (FCGRIIIA, EPHB1; **Fig. 5A**). Purified proteins were digested with trypsin or chymotrypsin, labelled with DEL, and mixed pairwise, before mass spectrometry analysis (**Fig. S3A**). Glycans in α-DG were detected both in the glycoproteome data and in the model protein data for HEK293^WT^ and HEK293^SC^, but not in HEK293^nO-Man^, in agreement with previous data (**Supplementary material**). The β-DG glycosylation identified in glycoproteome data was not identified on the reporter protein (**Supplementary material**). Analysis of recombinantly-expressed CADH1 ectodomain yielded similar data to our previous studies (17) and glycoproteome data in this study, confirming that O-Man is present in HEK293^WT^ and HEK293^SC^, but not in HEK293^nO-Man^. We then sought to express CDH11 to shed light on the unique glycosylation patterns identified on this protein in this and previous datasets. Soluble ectodomain of CDH11 presented glycans in HEK293^WT^ and HEK293^SC^, but not in HEK293^nO-Man^. We also identified O-GalNAc at the same sites for O-Man, suggesting that absence of O-Man enzymes allows O-GalNAc glycosylation in Golgi apparatus (**Supplementary material**). We then focused on the non-canonical substrates, none of which showed O-Man glycosylation in either HEK293^SC^ or HEK293^nO-Man^ cells (**Supplementary material**). This was surprising since O-Man was identified on ITGB1, ITGA5, ITGAV, F11R earlier in this study and in our previous studies (18). We further investigated these findings by expressing DAG1 in its native transmembrane form, to assess glycan pattern both on α-DG and β-DG. We therefore generated a full length DAG1 construct with 3xFLAG tag at the C-terminal end and stably overexpressed it in isogenic HEK293^WT^, HEK293^SC^ and HEK293^nO-Man^ cells using Zinc Finger Nucleases KI on AAVS1 Safe Harbor locus (**Fig. S4A**). Expression of the construct was confirmed by means of Western Blotting (**Fig. S4B**). The expressed transmembrane protein was purified (**Fig. S4B**) using M2-conjugated magnetic beads (anti-FLAG mouse IgG) and subjected to proteolytic digestion on S-trap, diethyl-labeling and MS analysis (**Fig. S3B**). We confirmed O-Man glycosylation was on α-DG but not on β-DG, indicating that the O-Man glycosylation on β-DG, and other non-canonical O-Man substrates, has low occupancy (**Supplementary material**). Taken together, these results demonstrate that O-Man glycosylation may occur on a wider range of substrates, albeit at low stoichiometry.

**Figure 5.**
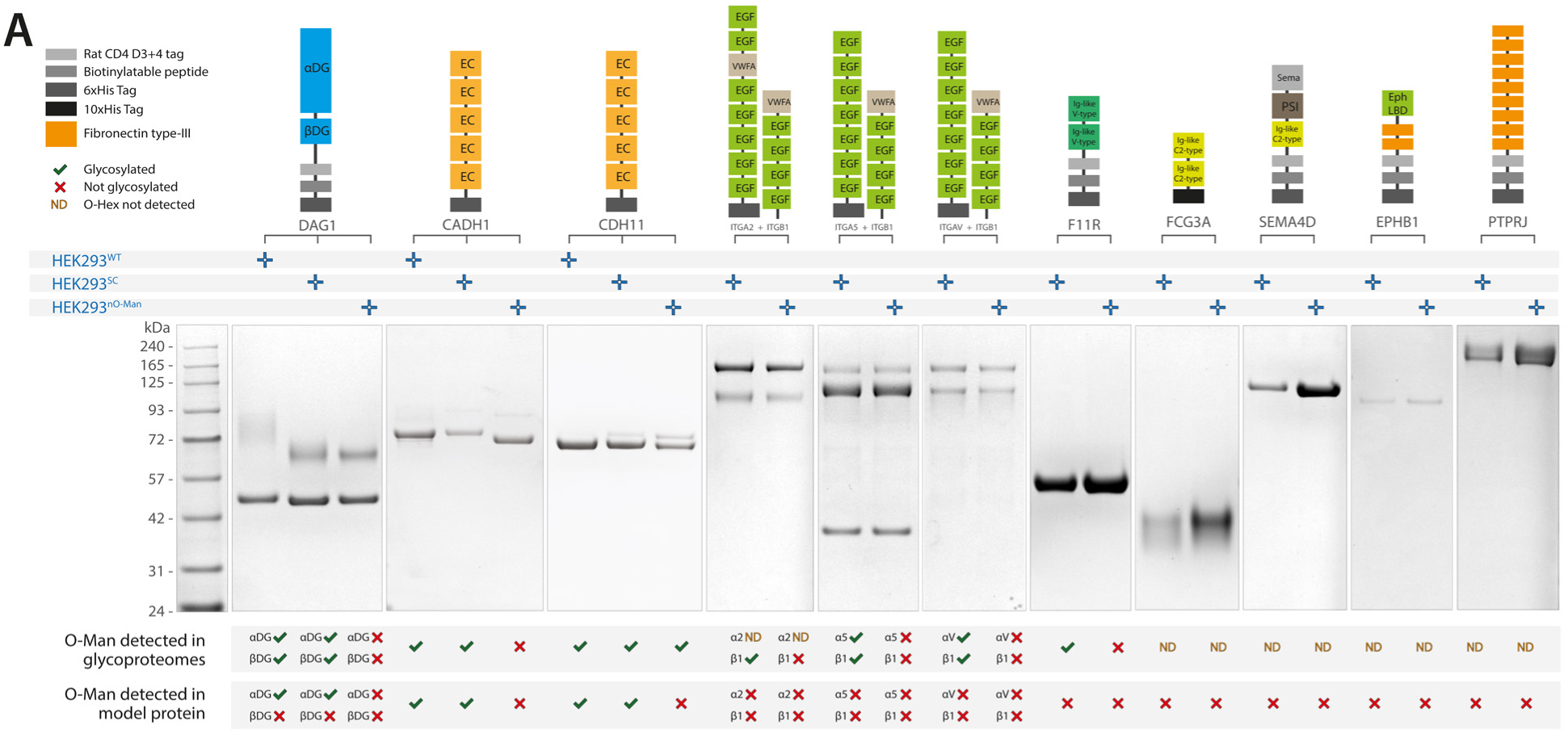
Analysis of O-Man using recombinantly expressed reporters. **(A)** Overview of constructs design and proteins expressed in cells with different genetic backgrounds. From top to bottom are the design of the constructs with the protein expressed and relative protein domains (according to Uniprot information), the information on which cell line the constructs were expressed in, a representative Coomassie staining for expressed proteins (relative expression levels between different genetic backgrounds are not quantitative), a graphical report on glycosylation status of the proteins according to glycoproteomics data from crude membrane preparations and purified proteins (bottom).

### BC2L-A lectin and membrane glycoproteomics expand the C-Man glycoproteome

Targeted membrane glycoproteomics workflow used here not only led to the identification of new O-Man proteins, but also expanded the knowledge of the C-Man glycoproteome, due to the ability of the lectin BC2L-A to capture C-Man peptides and proteins (35). Collectively, this study identified 73 human and 2 bovine C-Man proteins (most likely from cell culturing media) across 12 datasets and on 17 protein domains. Several protein domains are targeted by C-Man glycosylation, with the overrepresentation of TSP type-1 domains as primary target for C-Man glycosylation. Interestingly, we identify known O-Man proteins (α-DG, plexins, protocadherins). Reassuringly, the genetic engineering for O-Man enzymes did not impact their C-Man glycosylation, in line with the specificity of DPY19 enzymes (44). Nevertheless, these results should be interpreted with care before the occupancy and biological relevance has been determined.

## Discussion

O-Man glycosylation has emerged as a widespread modification among specific adhesion molecules and receptors of eukaryotic cells. The discovery of three independent biosynthetic pathways targeting distinct proteins at the cell surface, including *α*-DG, cadherins and plexins, clearly points to important biological roles for O-Man in human physiology (20), and this notion is further supported by the diverse and severe developmental phenotypes in patients with defects in O-Man biosynthesis (27,28,45,46).

Here, we provide further expansion of the O-Man glycoproteome in five human cell lines and map pathway-specific biosynthesis using a panel of WT and isogenic HEK293 cells with combinatorial KOs of O-Man initiation pathways. This study improves the glycoproteomic workflow for O-Man analyses, allowing us to identify a total of 180 proteins as targets for human O-Man enzymes, and unveils details on substrate specificities at protein-, domain-, and site-specific levels for each O-Man glycosyltransferase family, thus advancing the knowledge on how cells orchestrate and fine-tune biosynthetic processes relevant to O-Man glycosylation (**Fig. S5-S6**).

The first analysis of the mammalian O-Man glycoproteome relied on the combination of genetic engineering of the glycosylation capacities and lectin weak affinity chromatography (LWAC) followed by quantitative O-glycoproteomics (13,19,40,47). The current workflow introduces key improvements for sensitive identification and quantification of O-Man glycosylations, including the use of a small, dimeric BC2L-A lectin for C- and O-Man glycopeptide enrichment and substitution of deuterated dimethyl for ^13^C_2_-diethyl stable isotope labeling for improved glycopeptide retention on C18 reversed phase chromatography, ionization, and concurrent elimination of “deuterium effect” for improved quantification (39). The genetic engineering for deconstruction of biosynthetic pathways combined with our glycoproteomic strategy reveals two major types of transmembrane proteins as acceptor substrates for O-Man glycosylation. The first type of protein substrates are characterized by unstructured, mucin-like domains, including *α*-DG, KIAA1549 and SUCO, which are densely O-Man glycosylated by POMT1/POMT2 enzymes in the ER before continued biosynthesis and elongation in the secretory pathway resulting in capped and complex O-Man glycans known as core-M1, -M2 and -M3/matriglycan (15). The second type of protein substrates, which includes cadherins, protocadherins, plexins, cMET and RON receptors, are characterized by O-Man glycosylation on distinct Ig-like folds. This study confirms and further expands the knowledge on substrate specificities of the TMTC1-4 and TMEM260 enzyme families, which have unique functions for O-Man glycosylation of EC- and IPT-domains, respectively. Although both TMTC1-4 and TMEM260 enzyme families reside and perform their glycosylation function in the ER, O-Man on EC- and IPT-domains does not undergo further biosynthetic elongation into complex structures, as shown previously (17,18,48) and in this study, using secreted reporter proteins (**Fig. 5, Fig. S3A**). Alpha-linked O-Man monosaccharides are substrates for POMGNT1 and POMGNT2, which catalyze the second biosynthetic step and addition of GlcNAc-β1-2 or GlcNAc-β1-4-Man-*O*-Ser/Thr structures, respectively (15). It is thus surprising that EC- and IPT-domains, akin to C-Man modified TSP-domains (49), can traffic the secretory pathway and be presented at the cell surface (or secreted as reporter proteins) with O-Man monosaccharide modifications without further structural elongation and capping. Most likely, the polypeptide context to which O-Man is attached plays a decisive role for the following biosynthetic steps and impacts whether O-Man can serve as a substrate for additional glycosylation reactions. Complex O-Man glycans are found on unstructured, mucin-like regions of *α*-DG (19) as well as in the highly disordered extracellular domain of KIAA1549 (not shown), which indicates that O-Man located on disordered polypeptides is a preferable acceptor for the POMGNT1 enzyme. We have not observed complex O-Man glycans on the disordered SUCO protein, despite data mining using MS-Fragger (50) for open-search queries (not shown), which is reasonable considering that SUCO is an ER-localized protein that should not encounter the Golgi-resident POMGNT1 enzyme. In contrast, O-Man monosaccharides located on folded EC-and IPT-domains may interact with POMGNT1 during their traffic through the secretory pathway, but it is reasonable to believe that O-Man on folded Ig-like domain is an inaccessible substrate, potentially due to steric clashes between/within the active site of POMGNT1 and folded Ig-like domains.

In this study, we further confirm that POMT1/POMT2 are highly selective enzymes that target mucin-like domains of only a few protein substrates (*α*-DG, KIAA1549 and SUCO). The molecular basis for POMT1/POMT2 selectivity is puzzling considering that there are hundreds of disordered proteins with dense Ser/Thr content trafficking the secretory pathway, *e.g.* mucins, yet none of these have been reported to undergo O-Man glycosylation, and we find no evidence in this study to contradict this observation. A consensus sequon for O-Man glycosylation has not been identified, however, distinct polypeptide sequences termed *cis*-controlling peptidic elements have previously been suggested to function as recognition determinants for protein specific installation of e.g. polysialic acid, LacdiNAc, mannose-6-phosphate and POMT1/POMT2 driven O-Man glycosylation (51). Future structural studies focused on the interaction between POMT1/POMT2 MIR-domains and *α*-DG, KIAA1549 ectodomain and SUCO substrates may provide further insight to unravel the molecular details of POMT1/POMT2 substrate selectivity.

Our differential analyses of cells with genetically deconstructed biosynthetic pathways further highlights that TMTC1-4 and TMEM260 have unique functions and dedicated roles for O-Man glycosylation of EC-and IPT-domains, respectively. EC-domains, commonly found on the ∼120 different members of the cadherin superfamily of adhesion molecules, are modified on two distinct β-strands (B and G) with O-Man projecting in opposite orientations perpendicular to the plane of each EC-domain, indicating that TMTC1-4 may regulate or fine-tune cadherin functions and binding properties, especially for the clustered protocadherins which are known to utilize the EC1-EC4 domain interface for homophilic trans-interactions (52). The TMTC1-4 enzymes share a common N-terminal architecture including the first seven transmembrane (TM) that constitute the *conserved GT-C module* (7). Conserved acidic residues located within the first ER-luminal loop connecting TM1 and TM2 are predicted to be important for catalytic functions based on homology and structural similarities with *e.g.* yeast PMTs, POMTs, TMEM260 and other GT-C enzymes (18,53). The *variable GT-C module* comprises the C-terminal ER-luminal domains that distinguish TMTC1-4 from other GT-C enzymes based on a variable number of tetratricopeptide (TPR) repeats in each isoenzyme (**Fig. 7**). TPR repeats are evolutionarily conserved structural motifs that facilitate various molecular interactions between proteins, domains, and short polypeptides (54), which may explain TMTC1-4 specificity, *i.e.* cadherin recruitment through EC-domain recognition by the TPR domains before O-Man glycans are added at the catalytic site. We believe that the same reasoning applies to TMEM260, which has an N-terminal *conserved GT-C module* responsible for catalytic activity (18), while the ER-luminal *variable GT-C module* likely recruits substrates through specific interactions between TPR repeats and IPT-domains. Notably, IPT-domains are modified on B-strands, which is required for receptor maturation and traffic to the plasma membrane but may also influence *cis*-interactions and receptor dimerization events at the cell surface. For Ig-like folds including both EC- and IPT-domains, it however remains unknown whether O-Man glycans are added to unfolded, partially folded, or completely folded domains, and further studies are necessary to resolve the molecular details of TMTC1-4 and TMEM260 interactions with their substrates, potentially through cryoEM or structural analysis in simplified systems using recombinantly expressed Ig-like domains and TPR repeats.

**Figure 6.**
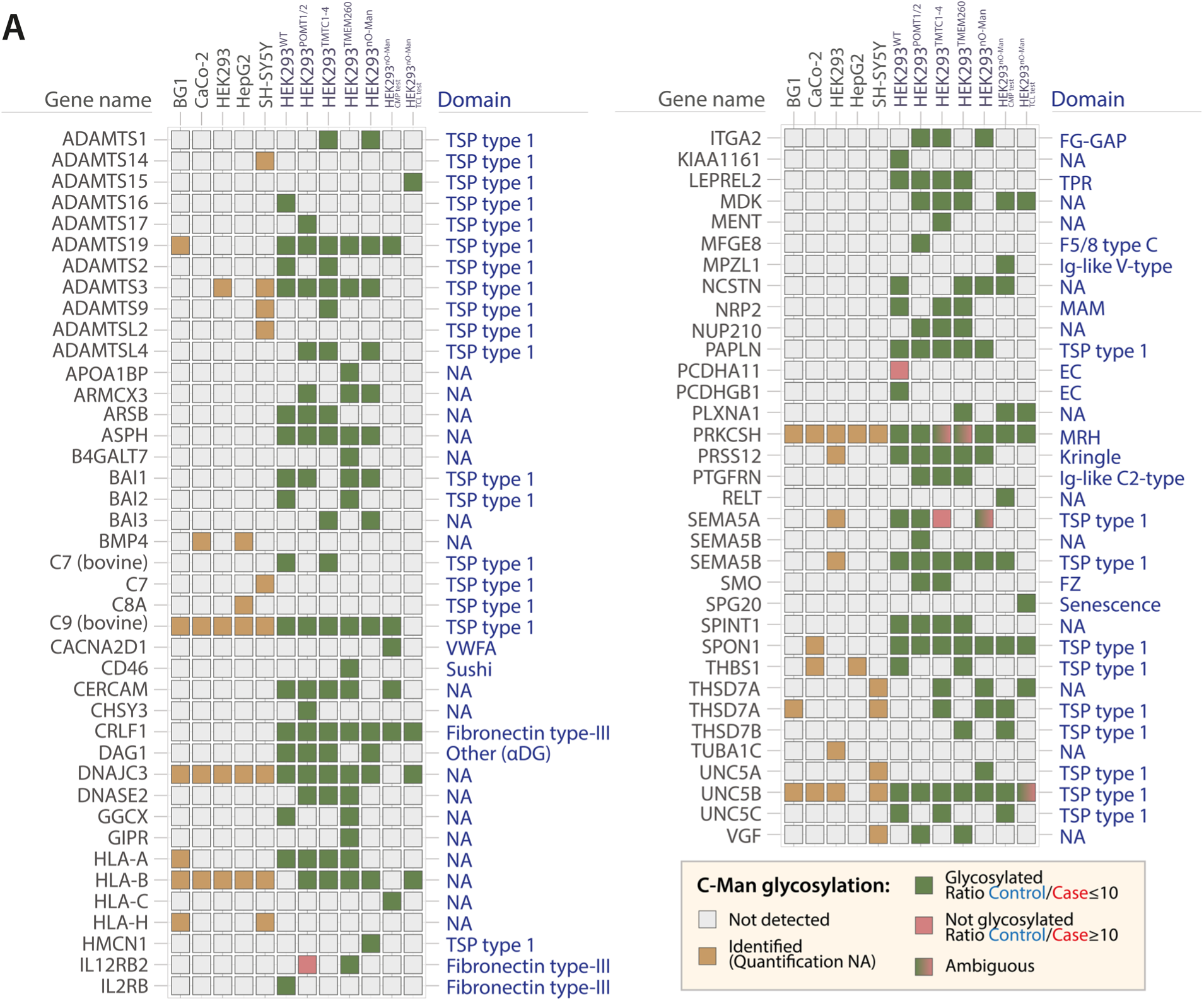
C-Man proteins identified across WT and glycoengineered human cell lines. **(A)** Overview of C-Man proteins identified in the datasets identified by BC2L-A lectin enrichment. Protein domain annotations were compiled based on Uniprot (May 2022). For proteins with same gene name but corresponding to multiple Uniprot accession numbers (e.g., HLA proteins), only one single gene name is reported.

**Figure 7.**
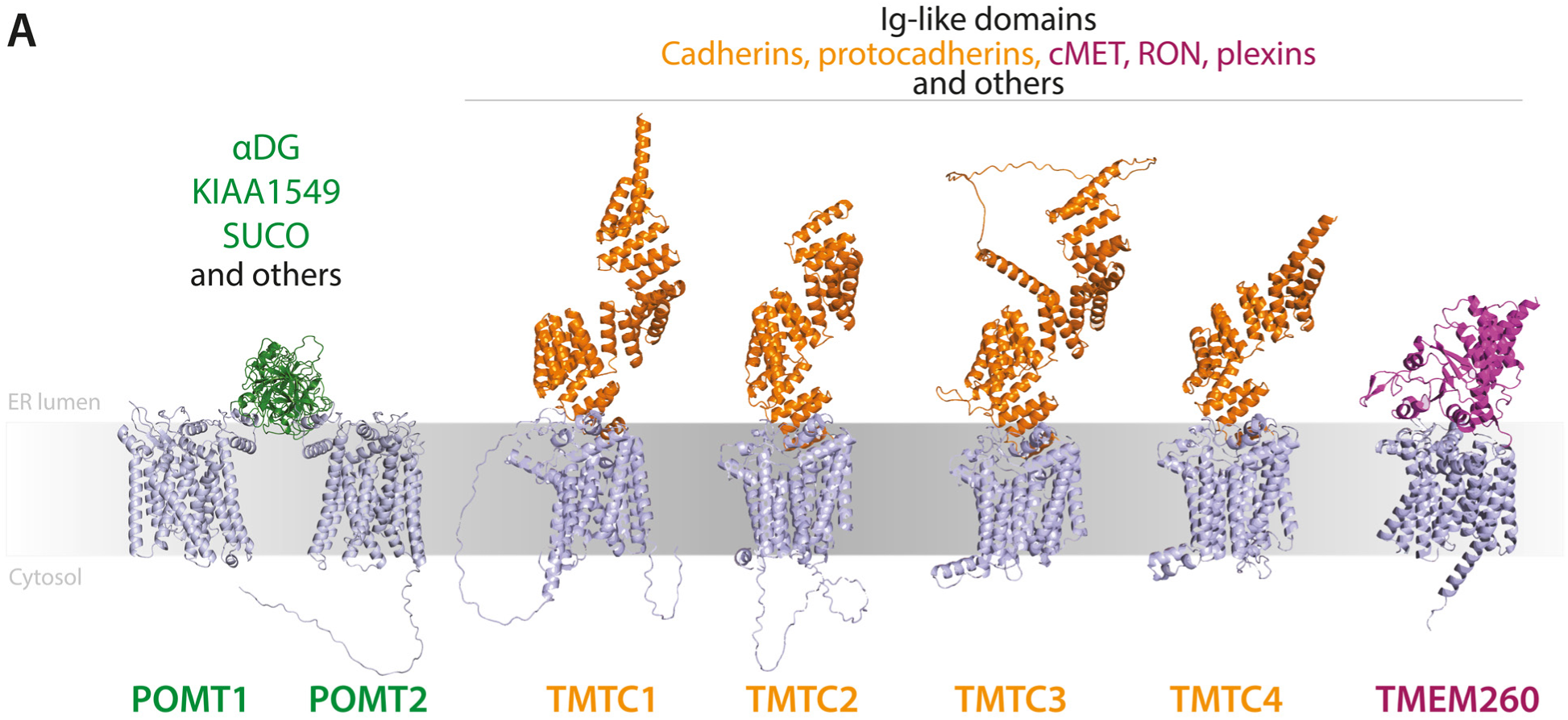
Three enzyme families for O-Man initiation in humans. **(A)** O-Man glycosylation is driven by three biosynthetic pathways: POMT1/2, TMTC1-4 and TMEM260. Each of these have specificities for canonical substrates and other substrates. Figures made using AlphaFold (68) and alignments made using PyMol (The PyMOL Molecular Graphics System, Version 2.5.2, Schrödinger, LLC.).

We also observe that all three biosynthetic pathways are capable of O-Man glycosylation on non-canonical protein substrates that do not fit the categorization described above. For example, protein disulfide isomerases (PDIA3, PDIA4 and PDIA6) are O-Man glycosylated by the TMTC1-4 family, even though they lack Ig-like domains (**Fig. 4A-B**). Previous studies have shown that TMTC3 may interact with PDIA3 (55), and it is possible that PDIs and TMTC1-4 isoforms form higher order molecular assemblies in the ER lumen, thus allowing PDIs to become O-Man glycosylated due to their close proximity to the catalytic site of TMTCs. Proximity-based glycosylation, unlike reactions driven by the intrinsic specificities of glycosyltransferases, may also explain O-Man on POMT1 and POMT2 enzymes in HEK293^POMT1/2^ cells, which are known to form heterodimers in the ER (56), or O-Man on TMTC1 and TMTC2 in HEK293^TMTC1-4^ cells, which indirectly suggests that functional TMTCs assemble as dimers in the ER.

Our results further showed that a subset of non-canonical substrates including β-DG, ITGA5, ITGAV, ITGB1 and F11R (**Supplementary material**), are O-Man glycosylated on β-strands of *e.g.* Ig-like C- or V-type domains. To further validate these findings, we adopted a targeted approach using a panel of recombinantly expressed and purified reporter proteins, however, we could not confirm O-Man glycosylations in bottom-up analyses (**Fig. 5A**), indicating that the occupancy of O-Man on non-canonical substrates is low and only detectable following enrichment procedures (**Fig. 3B**). We speculate that C- and V-type domains, together with other subclasses of Ig-like folds (**Fig. 4B-C**), transiently interact with TPR domains due to their overall structural similarity to EC- and IPT-domains, thus allowing a sub-fraction of substrates to become O-Man glycosylated at low occupancy. Therefore, we advocate that O-Man, as well as other types of O-glycosylations identified by large scale glycoproteomic analyses of complex samples, should be interpreted with care, and that such finding are validated by *e.g.* targeted analyses of isolated protein substrates before any major conclusions are drawn with respect to biological and/or functional relevance. This is also true for a number of POMT1/POMT2 substrates identified in HEK293^POMT1/2^ cells *e.g.* APP, ASTN1, GLA, HEXA, HSPA5, HYOU1, and TMEM43, where further studies are warranted to validate enzyme specificities and O-Man functions for these substrates.

Finally, our genetic deconstruction strategy with complete KO of O-Man initiation pathways in HEK293^nO-Man^ cells corroborates previous studies (13,17,18) and aligns well with the data presented in this study, demonstrating that combinatorial KO of *POMT1*, *POMT2*, *TMTC1-4* and *TMEM260* abolishes O-Man glycosylation on >100 protein substrates including *α*-DG, KIAA1549, the classical cadherins, protocadherins and plexin receptors (**Fig. 4**). Surprisingly, we note that CDH11, HNRNPA2B1, MCFD2 and EGFL2 are identified as glycoproteins in our HEK293^nO-Man^ dataset (**Supplementary material**). While we cannot rule out that these proteins are modified by O-Man, or other glycans that match the 162.0528 amu mass increment corresponding to hexose (e.g. glucose or galactose), we find it unlikely that a GT-C enzyme is responsible for glycosylation of these protein substrates. Further investigation is needed to resolve the biosynthetic basis for these modifications, especially for CDH11 and MCFD2, which are consistently identified with O-Hex modifications in our genetically engineered cell lines.

In conclusion, the human O-Man glycoproteome now covers >240 glycoproteins (180 identified in this study) involved in various functional networks, including those associated with cell-ECM interactions, cell-cell adhesion and receptor signaling (**Fig. S7**). The O-Man glycans are predominantly found on transmembrane proteins, of which a disproportionally large fraction is modified on unique protein folds, primarily Ig-like domains. This study establishes a roadmap by identifying substrates and defining specificities for the three families (POMT1/POMT2, TMTC1- 4 and TMEM260) of biosynthetic enzymes, which now can be used to guide further functional studies on O-Man glycosylation in human health and disease.

## Experimental procedures

### Experimental Design and Statistical Rationale

This study is based on four data packages (A-D), reported in **Supplementary material**; dataset A, aimed at improving the preparative methodology for O-Man glycoproteomics, was collected from experiments in HEK293 WT cells or similar and/or re-analyzed data from our previous study (17) (**Fig. 1**). Dataset B, aimed at identifying O-Man glycoproteins across five WT cell lines (**Figs. 2, 6**), is based on light diethyl-labelled (DEL) samples analyzed independently from each other by shotgun O-Man glycoproteomics. Dataset C is based on differential glycoproteomic analyses (**Figs. 3, 4, 6**) of five individual datasets with light/heavy DEL. Unless otherwise specified, membrane preparations of glycoengineered cell lines were labelled with light isotopes and control cell lines HEK293 with KO of *COSMC* and *POMGNT1* genes (HEK293 WT^SC/SC^) were labelled heavy isotopes. Dataset D is based on analyses of purified transmembrane or soluble proteins overexpressed in WT or glycoengineered cells (**Fig. 5**), labelled with diethyl isotopes before bottom-up analyses. All peptide spectral matches (PSMs) were identified by Proteome Discoverer 1.4 using probability-based scoring (Sequest-HT, p<0.01) and manual inspection of glycopeptide MS/MS identifications and MS1-level quantifications to ensure the accuracy of the assignments. Unless otherwise stated (Methods section), datasets are based on single shotgun experiment, with one biological and one technical replicate.

### Mammalian cell culturing

HEK293 and Caco-2 cells were grown in DMEM culturing medium (Sigma) with 10% FBS (Gibco) and 1% GlutaMAX (Gibco). The same medium, with further addition of 1% MEM Non-essential Amino Acid Solution (Sigma) was used for culturing of HepG2 cells. BG1 cells were cultured in RPMI 1640 medium (Sigma) supplemented with 10% FBS (Gibco) and 1% GlutaMAX (Gibco). SH-SY5Y cells were cultured in 50:50 mixture of DMEM medium (Sigma) and RPMI 1640 (Sigma), supplemented with 10% FBS (Gibco) and 1% GlutaMAX (Gibco). For protein production, cells were handled as described below. An overview of cell lines generated and used in this work can be found in **Supplementary table 3**.

### Constructs, guides design and genetic engineering

Knock-Out (KO) cell lines were generated according to published protocols (57,58). Parental cell lines and guide RNA (gRNA) plasmids used in this study have been described previously (18,59). Parental HEK293 cells were co-transfected with 1.0 µg each of PBKS-Cas9-2A-eGFP plasmid (Addgene plasmid #68371) and plasmid encoding for gRNA, as in **Supplementary table 1** using Lipofectamine 3000 (Thermo Scientific), according to manufacturer’s protocol. Cells were FACS-enriched 48 hours later, following fluorescent signal and further single-cell sorted seven days thereafter. Indel Detection by Amplicon Analysis (IDAA) guided the selection of knock-out clones, using primers in **Supplementary table 2** (57,58) which were further validated by Sanger Sequencing. For model proteins in **Figure 5**, constructs for soluble DAG1 (Addgene plasmid #51651), ITGA2 (Addgene plasmid #51910), ITGB1 (Addgene plasmid #51920), ITGA5 (Addgene plasmid #51909), ITGAV (Addgene plasmid #51919), SEMA4D (Addgene plasmid #51827), EPHB1 (Addgene plasmid #51750) and PTPRJ (Addgene plasmid #51816), as well as the plasmid F11R pEBio (Addgene plasmid #61486) used for further cloning were a gift from Gavin Wright (60) and obtained through Addgene. For the generation of the data in this work, plasmid F11R pEBio (Addgene plasmid #61486) was modified by moving the 726bp region isolated by digestion, performed with NotI-HF® (NEB, cat. R3189) and AscI (NEB, cat. #R0558) according to manufacturer’s recommendations, onto the vector for PTPRJ (Addgene plasmid #51816), previously digested with the same enzymes, using T4 ligase (NEB, cat. #M0202) according to manufacturer’s protocol. Ligated product was transformed to Stellar competent cells (TaKaRa) and plated on Agar plates with Carbenicillin (100µg/mL) for selection. The final plasmid was confirmed by Sanger Sequencing. Plasmid for CADH1 were described previously (17). Plasmid for CDH11 was synthesized and cloned by GeneScript on EPB71 vector (Addgene plasmid #90018).

Plasmid for expression of full-length DAG1 with C-terminal 3xFLAG tag was generated by amplification of DAG1 cDNA sequence using primers in **Supplementary table 2** from reverse-transcribed total mRNA from HEK293 cells. mRNA extraction was performed with RNeasy Mini kit (QIAGEN) and reverse-transcription with High-Capacity cDNA Reverse Transcription kit (Applied Biosystems), both according to manufacturers’ protocols. PCR product was ligated with InFusion ligation enzyme (TaKaRa) on EPB71 backbone (Addgene plasmid #90018) previously linearized by PCR. Ligation products were plated on Kanamycin-containing Agar plates (50µg/mL) and final plasmids confirmed by Sanger Sequencing, highlighting that *DAG1* gene presents the natural variant S14W (Uniprot: VAR_024335, dbSNP: rs2131107). Stable expression of DAG1 full length construct was performed by Zinc Finger Nuclease KI on AAVS1 safe harbor locus as previously described (18). Briefly, parental HEK293 cells were co-transfected with 3µg of donor plasmid and 1.5µg each of ZH001C and ZH001D using Lipofectamine 3000 (Thermo Scientific), according to manufacturer’s protocol. eGFP- and Crymson-positive cells were FACS-enriched 48 hours after transfection and single-cell sorted seven days thereafter. KI clones were identified by Junction PCR with primers and protocols as performed previously (18), and validated by Western Blotting for 3xFLAG tag (Antibody M2, Sigma).

### SDS-PAGE and Western Blotting

For SDS-PAGE separation of proteins, samples were mixed with 4X NuPAGE™ LDS Sample Buffer (Invitrogen), reduced with 5mM dithiothreitol (DTT) at 60°C and alkylated with 10mM 2-iodoacetamide (IAA) at RT in darkness. Proteins are separated using pre-cast 10-well NuPAGE™ 4 to 12%, Bis-Tris, 1.0–1.5 mm, Mini Protein Gels (Invitrogen) using MES buffer. Prestained Protein Ladder – Broad molecular weight (10-245 kDa) (Abcam, ab116028) was used as molecular marker. Gels are stained with InstantBlue® Coomassie Protein Stain (ISB1L) (Abcam, ab119211) and destained with water. For Western Blotting, SDS-PAGE-resolved gels were transferred to ethanol-activated PVDF membrane using a Tris-glycine transfer buffer (25mM Tris pH 8.0 at RT, 192 mM glycine, 10% ethanol in water). Membranes were blocked for 30 minutes at RT with blocking milk, made with 5% skim-milk powder diluted in 20 mM Tris pH 7.4, 150 mM NaCl, 0.1% Tween-20 (TBS-T) and incubated for one hour at RT with Mouse anti-FLAG (M2) antibody-HRP conjugated (Sigma) diluted 1:4000 in blocking milk. After 3x washes in TBS-T (5 minutes each), membranes were developed by SuperSignal™ West Pico PLUS chemiluminescent substrate (Thermo) using ImageQuant™ LAS 4000 (GE Healthcare). Membranes were stripped with 1X ReBlot Plus Strong Antibody Stripping Solution (#2504, Merck Millipore), blocked for 30 minutes in blocking milk, and incubated overnight at 4°C with anti-β-actin (Santa Cruz) diluted 1:1000 in blocking milk. After 3x washes in TBS-T, a 1:4000 dilution of anti-mouse HRP-conjugated secondary antibody into blocking milk (DAKO) was applied to the membrane for 1h at RT. Membranes were then washed and imaged as above.

### Glycoproteomic samples preparations

For glycoproteomics analyses (optimized workflow), cells were cultured in 7xT175 flasks to subconfluence and harvested by scraping in cold PBS. Cells were permeabilized in 150 mM NaCl, 50 mM HEPES pH 7.4, 25 µg/ml Digitonin (Sigma) and membranes further solubilized in 150mM NaCl, 50mM HEPES pH 7.4, 1% NP-40 according to Holden et al., 2009 (36). Membrane proteins were precipitated with 4X volumes of acetone overnight at -20°C and solubilized by sonication in 1 mL of 1 mg/mL RapiGest (Waters) in 50 mM ammonium bicarbonate buffer. Samples were reduced (5mM DTT, 30 min at 70°C), alkylated (10mM IAA, 30 min at RT in darkness) and unreacted IAA quenched with additional 5 mM DTT. Protease digestion was performed overnight with 25 µg of trypsin modified, sequencing grade enzyme (Roche) per sample and tryptic peptides were desalted using C18 Sep-Pack Cartridges (Waters). Diethyl labelling of Speedvac-desiccated peptides was performed essentially as in Jung et al, 2019 and in Larsen et al. 2023 (18,39). For differential glycoproteomics, labelled peptides were mixed 1:1 according to ratiocheck analyses by mass spectrometry. Peptide mixtures were dried using Speedvac, before treatment with 10U EndoH (NEB) in Sodium Acetate buffer, pH 5.5 overnight, and subsequently with 15U PNGaseF (Sigma) in Tris buffer, pH 8.6 overnight at 37°C in constant agitation at 750 RPM.

### BC2L-A beads and column preparation

BC2L-A lectin was produced using a pRSET-A construct encoding the full-length BC2L-A with an N-terminal 6xHis tag (35). The construct was electroporated into Rosetta™ 2 *E. coli* competent bacteria (Novagen) and transformed cells were selected on agar plates using Carbenicillin (100µg/mL) and Chloramphenicol (34µg/mL). A 1L culture of selected bacteria (OD600∼0.7) was induced for 3 hours with 0.1 mM Isopropyl β-D-1-thiogalactopyranoside at 37°C. Bacteria were harvested, washed in PBS and lysed with TES buffer (200mM Tris-HCl, pH 8.0, 0.5 mM ethylenediaminetetraacetic acid (EDTA), pH 8.0, 500 mM Sucrose) supplemented with 1 mM phenylmethylsulfonyl fluoride and 3 µg/ml Pepstatin A. Lysate was clarified by centrifugation at 27000×*g* and supplemented with NaCl at final concentration of 150 mM. Ni-NTA Agarose (Qiagen) was added to the clarified lysate for 30 minutes and moved to stationary column. Agarose was washed with 6 column-volumes (CVs) of 20mM sodium phosphate, pH 8.0, 900mM NaCl, with 6 CVs of 20mM sodium phosphate, pH 8.0, 150 mM NaCl, 10mM imidazole, pH 8.0 and eluted with 20mM sodium phosphate, pH 8.0, 150 mM NaCl, 250 mM imidazole, pH 8.0. Purified lectin was assessed by SDS-PAGE, buffer exchanged to 50mM HEPES, 150 mM NaCl, 1mM CaCl_2_ using Zeba™ Spin Desalting Columns, 7K MWCO (Thermo Scientific), and conjugated to Pierce NHS-Activated Agarose Dry Resin (Thermo Scientific) according to manufacturer’s protocol. A 3.5m LWAC column was prepared as previously (18,35), equilibrated with 25 mM Tris pH 7.4 at 4°C, 150 mM NaCl, 0.1mM CaCl_2_ and maintained at 4°C.

### Lectin Weak Affinity Chromatography (LWAC)

Digested samples are diluted to 1 mL with BC2L-A running buffer (25mM Tris pH 7.4 at 4°C, 150 mM NaCl) containing 0.1mM CaCl_2_, filtered with 0.22µM PVDF filters before LWAC. A 3.5-meter column with BC2L-A agarose beads was used for chromatographic separation at 100 µL/min using Äkta purifier (GE Healthcare) at 4°C. BC2L-A LWAC column was washed with running buffer until UV signal (A210) was lower than 5 mAU. The elution of glycosylated peptides was achieved with a 20 mM EDTA solution in running buffer. Elution fractions were acidified by trifluoroacetic acid (TFA) and EDTA precipitated by freeze-thaw cycle and centrifugation. Before MS/MS analysis, samples were stage tip-desalted (Empore disk-C18, 3M).

### Expression and purification of model proteins

HEK293 adherent cell lines with selected genetic background were seeded on 10-cm dishes previously coated with poly-L-lysine (Sigma, 0.1mg/mL). 10µg of plasmid were transfected on each 10-cm dish using 30µg polyethyleneimine (PEI) MAX, both diluted in OptiMEM (Gibco). Media were changed the following day to F17 Expression Medium (Gibco) with 2% GlutaMax or OptiMEM (Gibco) and supernatants were harvested 5 days thereafter. Harvested media were filtered to remove particles and used for Ni-NTA purification. For CADH1 and CDH11, 0.5mL Ni-NTA Agarose slurry (QIAGEN) each 50mL of supernatant were equilibrated in 1X Binding buffer (20 mM Tris, pH 8.0, 500 mM NaCl, 3 mM CaCl_2_, 5 mM Imidazole), added to supernatants supplemented to a final concentration of 20 mM Tris, pH 8.0, 500 mM NaCl, 3 mM CaCl_2_, 5 mM Imidazole and incubated overnight at 4°C in constant rotation. Beads were then moved to column and washed once with 5 CVs of binding buffer, a similar volume of wash buffer (20 mM Tris, pH 8.0, 500 mM NaCl, 3 mM CaCl_2_, 20 mM Imidazole) and eluted with 2.5 CVs of elution buffer (20 mM Tris, pH 8.0, 500 mM NaCl, 3 mM CaCl_2_, 200 mM Imidazole). For the remaining proteins, 0.5mL Ni-NTA Agarose slurry (QIAGEN) for each 50mL of supernatant were equilibrated in 1X Binding buffer (50 mM Sodium Phosphate buffer pH 8.0 at RT, 300mM NaCl, 10 mM imidazole), added to supernatants supplemented to a final concentration of 50 mM Sodium Phosphate buffer pH 8.0 at RT, 300mM NaCl, 10 mM imidazole and incubated overnight at 4°C in constant rotation. Beads were moved to column and washed once with 5 CVs of wash buffer (50 mM Sodium Phosphate buffer pH 8.0 at RT, 500 mM NaCl, 20 mM imidazole) and eluted with 2.5 CVs of elution buffer (50 mM Sodium Phosphate buffer pH 8.0 at RT, 150 mM NaCl, 300 mM imidazole). Fractions were inspected by SDS-PAGE for purified proteins before further analyses. Samples were buffer exchanged with Zeba Spin columns (Thermo Fisher) to 5mM HEPES pH 7.4, 5mM NaCl and stored at -80°C. All proteins in **Fig. 5A** are expressed independently, except for ITGA2, ITGA5 and ITGAV which are co-expressed with ITGB1, as previously recommended in original study using these constructs (60). This was done by co-transfecting equal amounts of plasmids (5µg of each ITGA2, ITGA5 and ITGAV + 5µg ITGB1) in each plate and performing purification as above. Regarding full length DAG1, purification was performed using dynabeads conjugated in-house with anti-FLAG M2 monoclonal antibody (Sigma-Aldrich). Beads conjugation was performed with a procedure adapted from previous protocols (61). Briefly, anti-FLAG M2 monoclonal antibody (Sigma-Aldrich) was conjugated to Epoxy Dynabeads™ M-270 (InVitrogen) at a ratio of 20µg per 1mg beads in 0.1M Na-Phosphate buffer, pH 7.4 with gradual addition of ammonium sulfate to a final concentration of 1M. Conjugation was performed overnight at 30°C in gentle rotation. Beads were washed with PBS and PBS+0.5% Triton-X100 and stored in 10% glycerol, 0.02% NaN_3_ at a final concentration of 150 ug beads/µL. Conjugated beads were washed in borate buffer (pH 8.5) and crosslinked using a 20mM solution of dimethyl pimelimidate (Thermo Scientific) in borate buffer (pH 8.5) for 2 hours. Beads were washed with 100mM Tris pH 8.0 and PBS and used for affinity enrichment of DAG1. DAG1-expressing cells were lysed in a cell pellet:lysis buffer ratio of 1:2 using 50mM HEPES pH 7.4, 50mM KOAc, 2mM MgCl_2_, 300mM NaCl, 0.5% Triton X-100, 0.5% Tween-20. Input lysate was cleared with 21000×*g* centrifugation and incubated for 1h in rotation at 4°C with anti-FLAG beads, pre-cleared in lysis buffer. Beads were washed in lysis buffer lacking Triton and eluted with 10% SDS (55µL) for 20 minutes in agitation. Approximatively 10% of elution volume was used for WB analysis, 45% was used for SDS-PAGE analysis to confirm absence of contaminants and the remaining 45% for S-Trap digestion and proteomics analysis, as below.

### S-Trap digestion of model proteins

10µL of purified proteins were reduced with 10mM DTT and alkylated with 20mM IAA. Unreacted IAA was quenched with the addition of 10mM DTT, and SDS was added to a final concentration of 5%. Digestion was performed using Micro S-Trap columns (ProtiFi) following the manufacturer’s S-Trap micro high recovery quick card 2, using ammonium bicarbonate (AMBIC) buffer instead of TEAB. Digestion was performed overnight at 37°C using modified trypsin or chymotrypsin (only for FCGRIIIA). Eluted samples were desalted by in-house packed Stage tips (Empore disk-C18, 3M) before diethyl labelling. Labelled peptides from soluble constructs were mixed according to Nanodrop measurements (Absorbance = 205 nm) and desalted before mass spectrometry analysis. Peptides from full-length DAG1 purified from total cell lysates were labelled with DEL, desalted, and analyzed individually using mass spectrometry (**Figure S3)**.

### Mass spectrometry analyses

Samples were analyzed using mass spectrometry following protocols implemented in our previous work (18) with slight modifications. Peptide and glycopeptides were dissolved in 0.1% formic acid and injected using an EASY-nLC 1000 or -nLC 1200 (Thermo) system coupled to a Fusion Tribrid or Fusion Tribrid Lumos mass spectrometer (Thermo), respectively. The nLC systems utilized a single analytical column packed with Reprosil-Pure-AQ C18 phase for sample separation. For glycoproteome analyses, gradient elution was performed using solvent A (0.1% formic acid) and solvent B (acetonitrile with 0.1% formic acid), with a stepwise increase from 5% to 20% B over 95 minutes, followed by a gradient from 20% to 80% B over 10 minutes, and a final hold at 80% B for 15 minutes. For bottom-up analyses of S-Trap digested samples, a similar gradient elution was employed, starting from 2% to 25% B over 65 minutes, followed by a gradient from 25% to 80% B over 10 minutes, and a final hold at 80% B for 15 minutes. For ratiochecks, a similar gradient elution was employed, starting from 2% to 25% B over 95 minutes followed by a gradient from 25% to 80% B over 10 minutes, and a final hold at 80% B for 15 minutes. Mass spectrometry analysis included precursor MS1 scans (m/z 355 to 1,700) acquired in the mass spectrometer at a resolution of 120,000. Subsequently, Orbitrap Higher-energy dissociation (HCD)-MS/MS and electron-transfer/collision-induced dissociation (ETciD)-MS/MS were performed on multiply charged precursors (z = 2 to 6). Only HCD fragmentation was performed for ratiocheck samples. Data-dependent fragmentation events were triggered by a minimum MS1 signal threshold of 10,000 to 50,000 ions. The MS2 spectra were acquired at a resolution of 60,000 for both HCD and ETciD methods. The comparison of HEK293^SC/SC^ vs. HEK293^nO-Man^ was analyzed on Fusion Tribrid Lumos and Fusion Tribrid instruments.

### Mass spectrometry data search

Mass spectrometric data (.raw files) was processed by the Proteome Discoverer (PD) 1.4 software (Thermo Fisher Scientific) and the Sequest HT node. Individual .raw files from crude membrane preparations (Dataset A-C) were searched against the canonical human proteome downloaded (January 2013) from the UniProtKB database (http://www.uniprot.org/) (62). Dataset D from purified model proteins was searched against a FASTA file containing the sequences of all model proteins in this study (available in **Supplementary material**). Detailed information on .raw files and processing is available through Proteome Xchange database (see **Data availability** section). Parameters for data processing are summarized in **Supplementary table 4.** Fragment ion mass tolerance was set to 0.02 Da and up to n=2 missed trypsin cleavages (both full- and semi-specific) were allowed during each search. Peptide confidence levels were determined using the Target Decoy PSM Validator node, and only identifications with high confidence (P < 0.01) were considered. Filter was applied to retain only minimum Peptide Confidence = High, minimum Search Engine Rank = 1 and Peptide Mass Deviation = 7.0 ppm. Data from the comparison HEK293^SC^ vs. HEK293^nO-Man^ analysed with Fusion Tribrid alone were processed as above, for **Fig. 1D** and **Fig. 1E**, while data from both Fusion Tribrid Lumos and Fusion Tribrid instruments were analyzed jointly in Proteome Discoverer for **Figs. 3, 4, 6** and supplementary figures. All spectral matches were inspected and validated through manual examination and reported at this stage in **Supplementary material**.

### Mass spectrometry data processing and visualization

Manually inspected and validated spectral matches from glycoproteome data are exported from Proteome Discoverer and further processed in Microsoft Excel. To perform domain annotations for glycosylated sites, we relied on the “Domains” annotation provided by Uniprot (May 2022) for categorization. purposes with slight modifications for laminins, integrins and DAG1. Laminin domain annotated as “Domain I” and “Laminin Domain I” are both classified as “Laminin Domain I”; laminin “Domain II” is annotated as “Laminin Domain II”. The domain annotated as “Laminin Domain II and I” is reported separately in figures, but not accounted as separate individual domain (i.e., in domain accounting for instance in **Fig. S6**) this does not count as individual domain, and in putative specificity it follows the information obtained in PSMs from “Laminin Domain I” and “Laminin Domain II”). For integrins, domains with no annotated domain report annotation of heavy and light chain, according to Uniprot data. For DAG1, regions of the protein are reported according to Uniprot information and previous literature (13). Both integrin and DAG1 regions are accounted for in the count of individual domains (i.e., in **Fig. S6**). “Domain annotation not available” indicates a region of the protein where there is no annotation of a domain in Uniprot database. Gene names are annotated according to Proteome Discoverer output, with the only exception of the protein C14orf166B, reported with its updated (August 2023) gene name LRRC74A. For proteins with same gene name but corresponding to multiple Uniprot accession numbers (eg. HLA proteins), only one single gene name is reported according to Uniprot information (August 2023). Analogously, for Uniprot accession numbers corresponding to multiple gene names/isoforms (eg. HLA proteins), only the main isoform according to Uniprot database is reported. PSMs are classified as O- or C-Man according to “Modification” column. To ensure accuracy of reported data, all glycoPSMs with “W” were manually inspected to ensure the accuracy of site-specific assignments by HCD and/or ETciD, particularly for peptides with sequence containing Ser/Thr proximal to a Trp residue. The glycan type of glyco-PSMs with insufficient MS2-data to support site-assignment was categorized as “Ambiguous”. All inspected PSMs (including contaminant bovine proteins detected and with any value in the column “# Protein Groups”) are used for PSMs counts. A PSM with at least one of the two types of sugar linkages counts as “O- and C-Man glyco-PSM” and excluded from the count for the “O-Man glyco-PSM” and “C-Man glyco-PSM”. For protein counts, only proteins with “# Protein Groups” = 1 are counted, excluding ambiguously identified proteins. A protein is a C- or O-Man protein if it contains the respective type of sugar on at least one PSM from the inspected list. Proteins with at least one O-Man PSM and one C-Man PSM is considered “O-and C-Man protein” and excluded from the count for the “O-Man protein” and “C-Man protein”.

For Figures 2C, 4A-B and 6, O-Man and C-Man PSMs are isolated and presented separately. For this protein visualization, only proteins with “# Protein Groups” = 1 are considered, excluding ambiguously identified proteins. PSMs are categorized into ratio groups, where each PSM is assigned to only one ratio category. For each gene name, one entry is kept for each domain on which the sugar has been identified and on the ratio category using “Remove Duplicates” function in Excel. Proteins with more than one category of ratio for each domain are collapsed into one entry with category of ratio as a merge of the two (e.g. 10≤x<100 and x≥100 becomes x≥10). In case of two non-adjacent groups, the corresponding square of the heatmap will have double color (in the case of x≥100 and 0.1≤x<10). The same procedure has been done analogously to O- and to C-Man entries. A ratio difference of 10 has been used as threshold level for difference between case and control based on ratiocheck results (**Fig. S2**) and with calculations mentioned in Results section.

In Figure 2, specifically for the “Other domains” category, if a protein has glycosylation on more domains classified as “Other domains,” it is represented graphically only once. Conversely, in Figure 4, to depict the specificity of each enzyme family on different domains, the protein is represented multiple times corresponding to the number of domains within each respective category of the heatmap.

For ratiocheck data processing and visualization, only PSMs with Quant Usage = “Used” are considered in the calculations. For calculation purposes, Ratios that equal to 0 are substituted with 0.00001.

Data and figures for heatmaps, barplots, ratiocheck distribution histogram, upset plot, lollipop plot were processed and generated using R, with the packages ggplot2 and upsetR, among the others, and further modified using Adobe Illustrator. Data intersections for Venn diagrams and barplots were performed using the online tool InteractiVenn (63) and further processed offline.

Gene ontology enrichment analyses were performed using ShinyGO 0.77 (64), using human pathway database GO Biological Process. No background list was uploaded; by ShinyGO default, the gene list was therefore compared with all protein-coding genes in the genome as background. FDR cutoff set at 0,05, “# pathways to show” set to 10, pathway size between 2 and 2000, removing redundancy and abbreviating pathways. Graphs were made on the same online tool by sorting pathway by fold enrichment (x-axis), coloring by -log10(FDR) and using the gene number as size of the terminal circles.

Figure S1 was made using data from studies in references (13,17–19,35). The 175 unambiguous and unique proteins identified were compiled after manual inspection following parameters in this method section. Note that protein with Accession Number P01889 reported as O-Man protein in (35) has been excluded for these criteria. List of proteins with transmembrane region was acquired in August 2023 via UniProt using the filters (taxonomy_id:9606) AND (keyword:KW-0812) AND (reviewed:true). Proteins with Signal Peptide were acquired likewise with the filters (taxonomy_id:9606) AND (keyword:KW-0732) AND (reviewed:true).

Figure S7 was made using STRING (65). All 241 O-Man proteins unambiguously (and manually inspected, following the parameters from this paragraph) identified in this study and in previous studies (13,17–19,35) were searched using STRING online tool against human proteome. Note that for this visualization, the data from the test experiment between CMP and TCL (Fig. 1B) were included. The full STRING network is visualized, with network clustered using k-means clustering (8 clusters) in STRING and leaving all the parameters as default. Colouring is made in STRING; proteins with SMART domain identifier SM00112 (Cadherin repeats) are coloured in red and with SMART domain identifier SM00429 (Ig-like, plexins, transcription factors) are in blue. The image was further modified using Adobe Illustrator. Two additional accession numbers reported in previous studies but not identified in STRING databases are manually added at the bottom of the figure.

## Abbreviations

amu: Atomic mass unit
BC2L-A: Burkholderia cenocepacia lectin A
C18: C18 silica column
Cas9: CRISPR associated protein 9
CAZy: Carbohydrate-Active enZYmes
C-Man: Protein C-Mannosylation
CMP: Crude Membrane Preparation
ConA: Concanavalin A
COSMC: C1GALT1 specific chaperone 1
DEL: Diethyl Label
DGC: Dystrophin-associated glycoprotein complex
DMEM: Dulbecco’s Modified Eagle’s Medium
DML: Dimethyl Label
Dol-P-Man: Dolichol phosphate mannose
EC: Extracellular Cadherin
ECM: Extracellular Matrix
eGFP: Enhanced green fluorescent protein
EndoH: Endoglycosidase H
ER: Endoplasmic Reticulum
ETciD: Electron-transfer/collision-induced dissociation
FDR: False Discovery Rate
gRNA: Guide RNA
GT-C_A_: Glycosyltransferase of the C superfamily and A class
HCD: Higher-energy collisional dissociation
HEK293: Human embryonic kidney 293 cells
Hex: Hexose
HexNAc: N-Acetylhexosamine
IDAA: Indel Detectionby Amplicon Analysis
Ig: Immunoglobulin
IPT: Immunoglobulin-like, plexins, transcription factors
KI: Knock-in
KO: Knock-out
LWAC: Lectin Weak Affinity Chromatography
MET/cMET: Hepatocyte growth factor receptor (HGF receptor)
MIR: Protein Mannosyltransferase, Inositol 1,4,5-trisphosphate receptor (IP3R) and
(RyR) [domain]: Ryanodine receptor
MS: Mass Spectrometry
MST1R/RON: Macrophage-stimulating protein receptor/Recepteur d’Origine Nantais
nLC: Nano-liquid chromatography
NP-40: Tergitol-type NP-40, Nonyl phenoxypolyethoxylethanol
O-Man: O-linked mannose-type glycosylation, O-mannosylation
PMT: Dolichyl-phosphate-mannose--protein mannosyltransferase
PNGaseF: Peptide-N-Glycosidase F
POMGNT1: Protein O-Linked Mannose N-Acetylglucosaminyltransferase 1
POMT1: Protein O-Mannosyltransferase 1
POMT2: Protein O-Mannosyltransferase 2
PSM: Peptide spectral match
PTM: Post-translational modification
RT: Room Temperature
SHDRA: Structural heart defects and renal anomalies (syndrome)
TCL: Total Cell Lysate
TMEM260: Protein O-mannosyl-transferase TMEM260
TMTC1: Protein O-mannosyl-transferase TMTC1
TMTC2: Protein O-mannosyl-transferase TMTC2
TMTC3: Protein O-mannosyl-transferase TMTC3
TMTC4: Protein O-mannosyl-transferase TMTC4
TSP type-1: Thrombospondin type-1
U: Unit
α-DG: Alpha-dystroglycan
β-DG: Beta-dystroglycan

## Acknowledgments

We thank colleagues at Copenhagen Center for Glycomics (CCG) for helpful discussions, Hans Bakker (Hanover Medical School, Germany) for sharing the BC2L-A lectin expression construct and Zhang Yang and Julie Van Coillie (CCG) for sharing the FCGRIIIA plasmid. This work was supported by the Danish National Research Foundation Grant DNRF107, the Mizutani Foundation for Glycoscience and a research grant (00025438) from VILLUM FONDEN.

## Data availability

The mass spectrometry proteomics data (.raw and .msf) have been deposited to the ProteomeXchange Consortium (66) via the PRIDE (67) partner repository with the dataset identifier PXD045597. The deposited data may be accessed using the following credentials: username; password: UrKmMHdB. This article contains supplemental data, including all validated data reported in tabular format. Annotated data used for generation of figures, as well as all codes utilized for generating figures and processing data are available at request to the corresponding author.

## Supplementary figures

**Figure S1.**
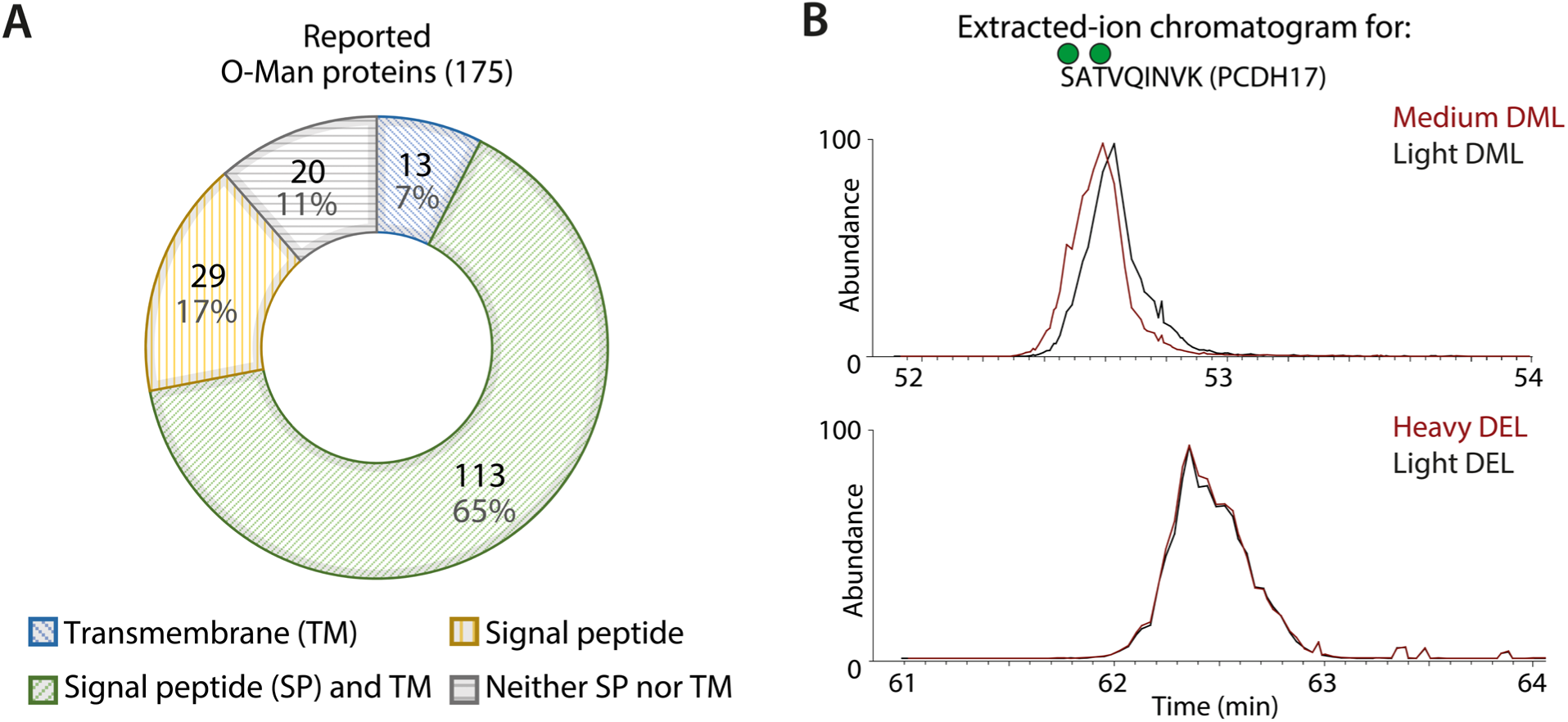
Data overview from O-Man glycoproteomic studies and comparison between labelling strategies. **(A)** 72% of O-Man proteins reported to date are transmembrane proteins and 17% have a signal peptide while only 11% have none of the two, suggesting that most human O-Man proteins are secreted or reside in the secretory pathway. The 175 unique O-Man proteins were extracted from datasets in (13,17–19,35), according to Methods section. **(B)** Nano-LC elution profiles of an O-Man glycopeptide labelled with heavy and light DEL (overlapping) and DML (displaced due to deuterium effect).

**Figure S2.**
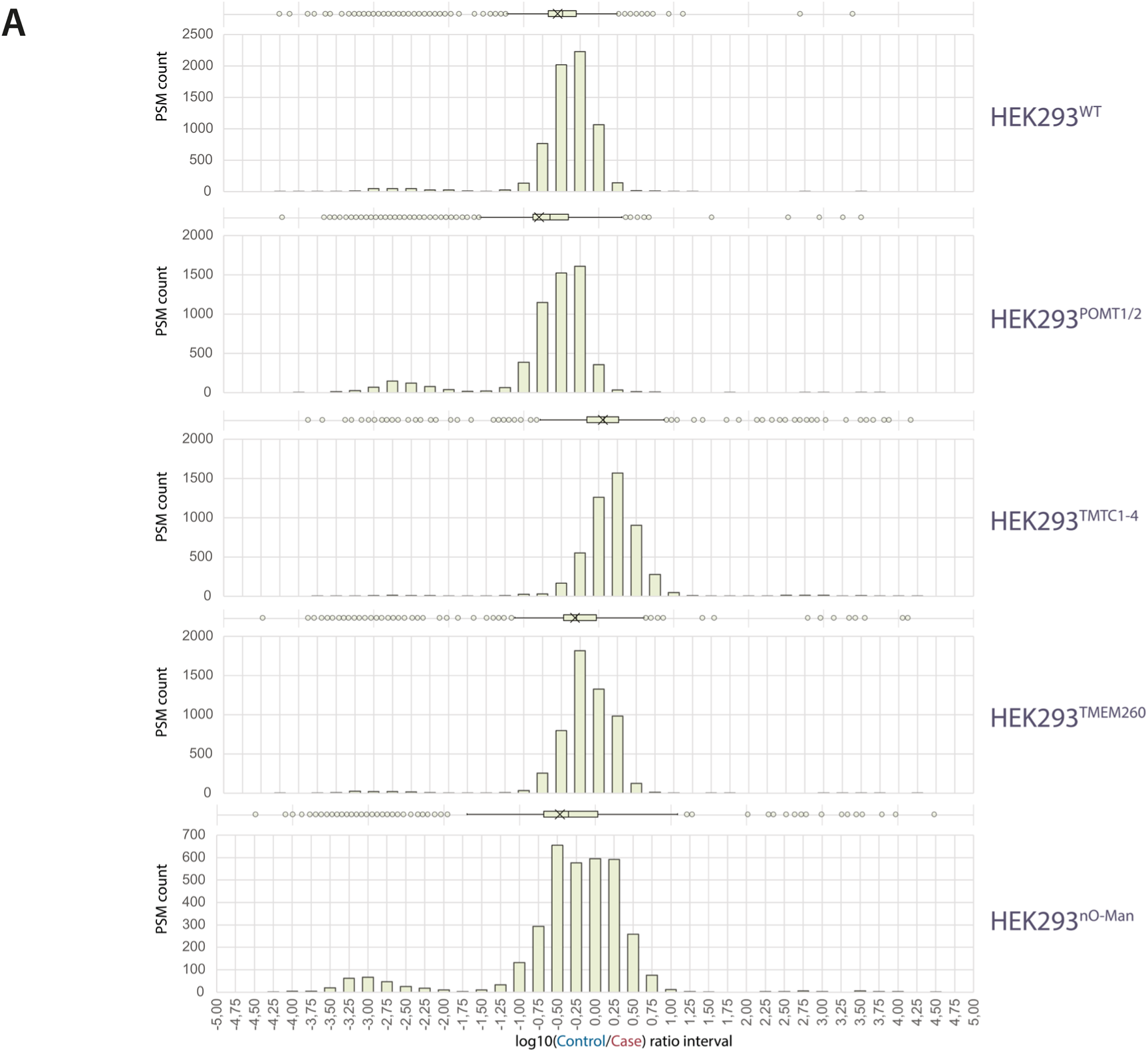
Ratiochecks for differential glycoproteomics samples. **(A)** Equal volume of samples labelled with heavy and light DEL were mixed and subjected to proteomics analysis. Calculated ratios (bins) from -5 (minimum value allowed, with log_10_ ratio = 10^-5^) to +5 (maximum value allowed, with log_10_ ratio = 10^5^). “PSM count” represent the frequency of Peptide-Spectrum Matches (PSMs) in each bin. Inter quartile range (IQR) is shown on top of histograms, where “x” depicts the median. The threshold interval for equal abundance of the two channels was set to 10-fold change. Cell line names (right) refer to the nomenclature in Figure 3.

**Figure S3.**
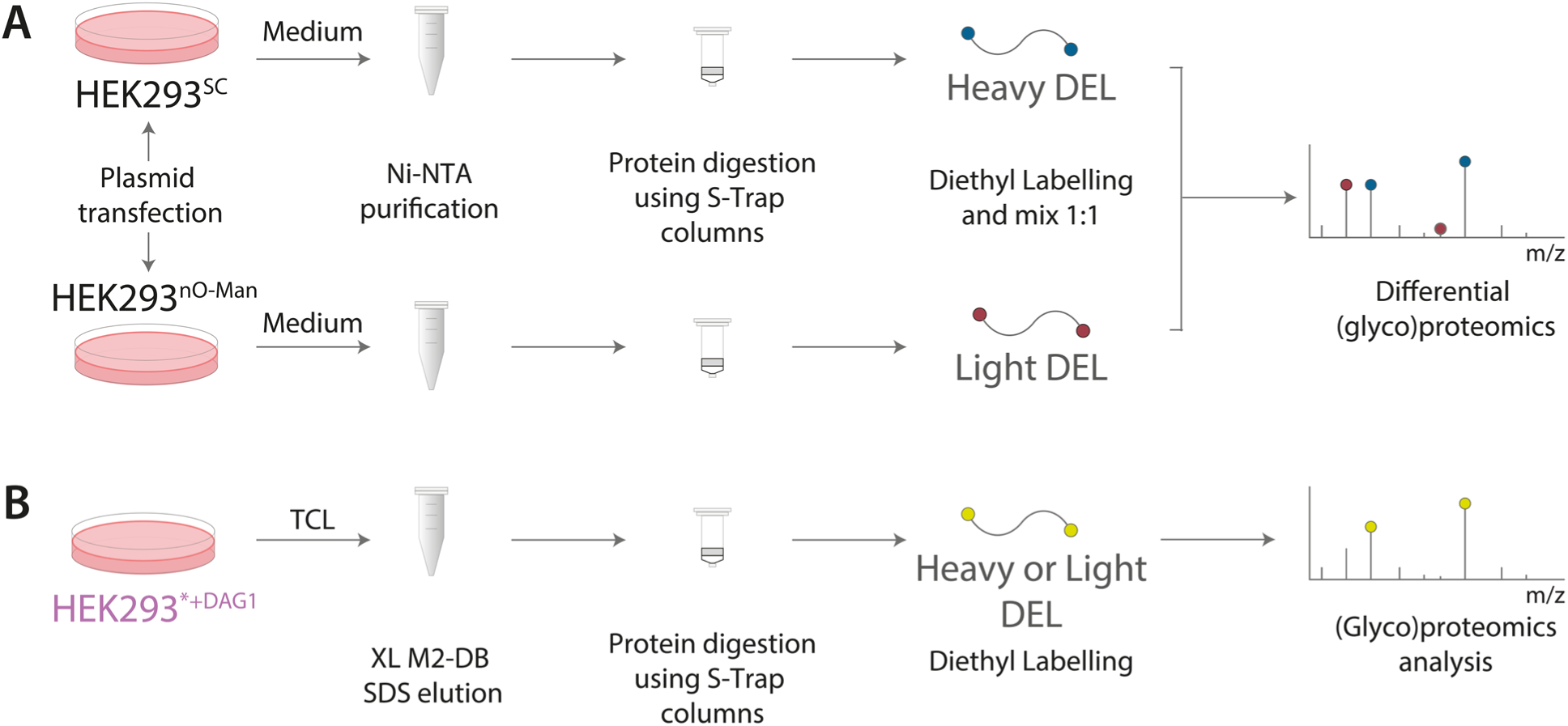
Schematic representation of workflow for (glyco)proteomics of reporters. **(A)** For secreted reporter proteins, glycoengineered cells are transiently transfected with plasmids and media are recovered after 5 days. Ni-NTA purification allows capture of model proteins which are then digested using S-Trap microcolumns and proteases. Peptides are labelled with DEL, mixed 1:1 according to nanodrop measurements (Absorbance = 205 nm) and subjected to MS analysis. **(B)** For full length DAG1, cell pellets from glycoengineered cells stably expressing DAG1-3xFLAG construct are lysed and subjected to capture using M2 antibody conjugated to magnetic beads (XL M2-DB). Proteins are diluted in SDS and digested with modified trypsin using S-Trap microcolumns. Peptides are labelled with DEL, desalted and subjected to MS analysis. Genetic backgrounds for the cells used here (in this figure, cumulatively referred to with a star*) are found in Figure S4.

**Figure S4.**
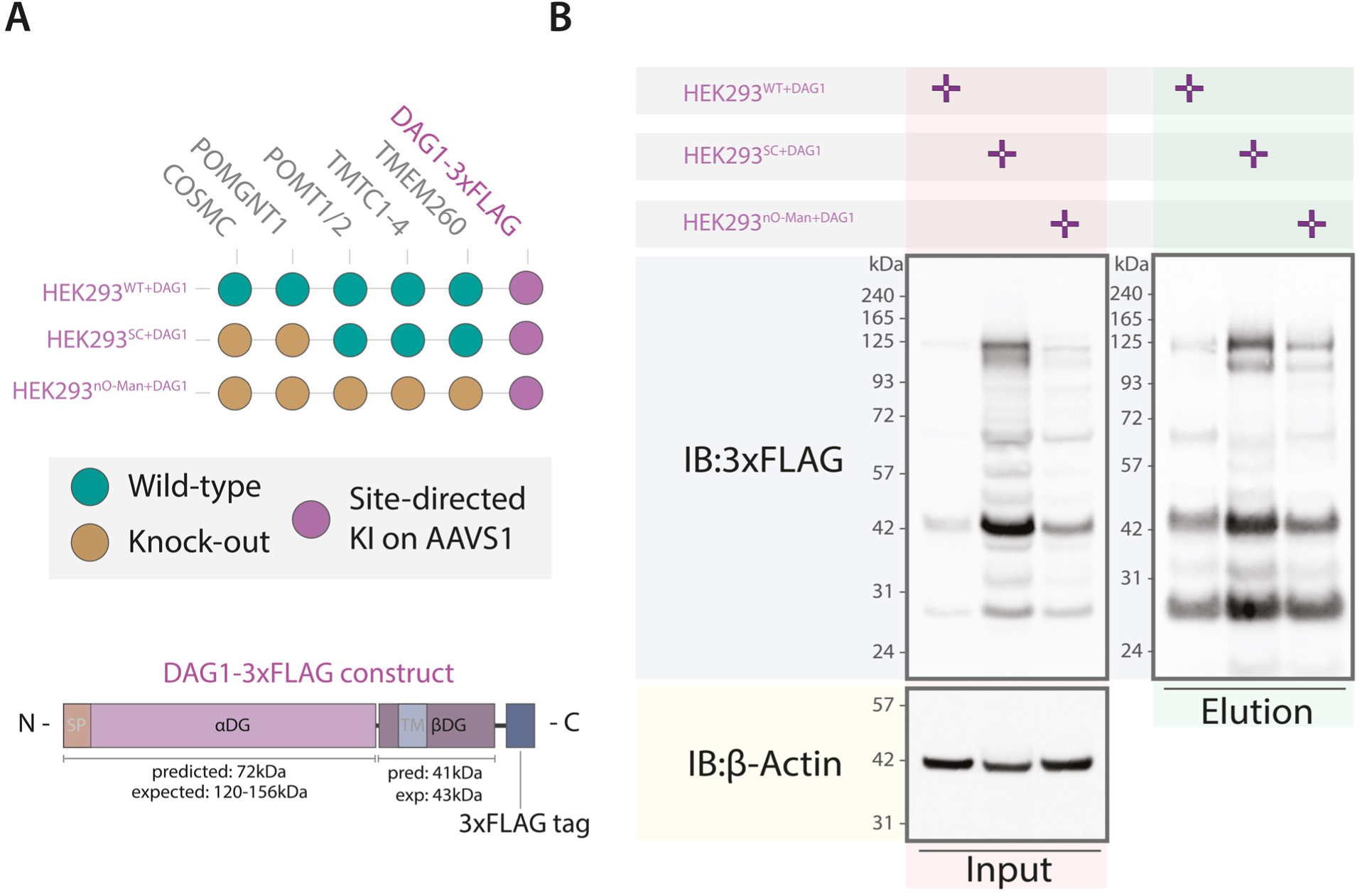
Generation and validation of glycoengineered cells expressing full length DAG1. **(A)** Overview of DAG1-3xFLAG construct and cell lines generated with KI on AAVS1 locus using zinc finger nucleases. **(B)** Western blot for Input and Elution from anti-FLAG resin confirms expression and enrichment of DAG1 used for proteomics studies as above. Sizes are reported as in (23). The established cell lines show that DAG1-3xFLAG is expressed at different levels.

**Figure S5.**
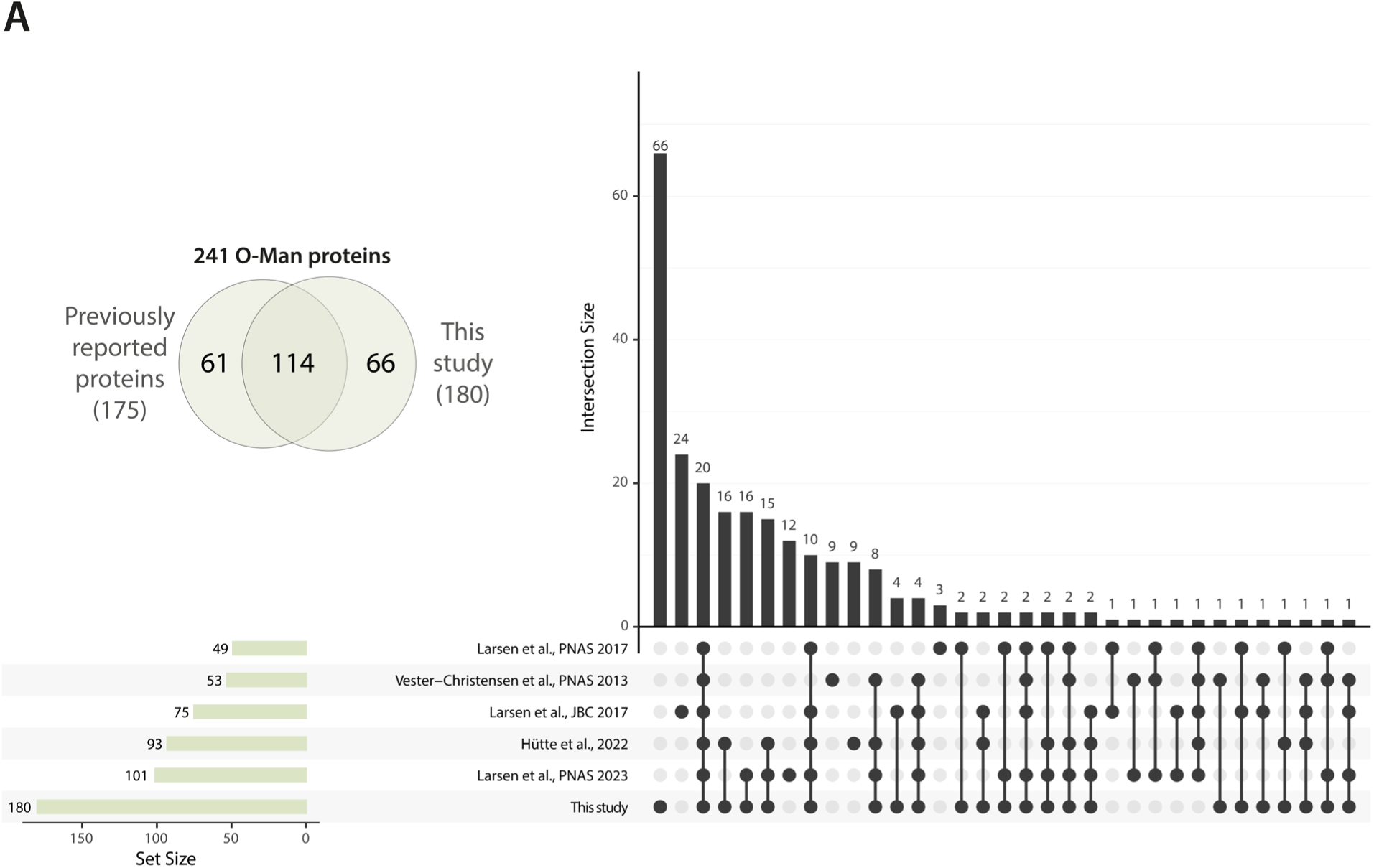
Comparison of O-Man proteins identified in this study and previous glycoproteomics studies. **(A)** Of the 180 proteins identified in this study, 66 were not reported in previous glycoproteomics studies (13,17–19,35).

**Figure S6.**
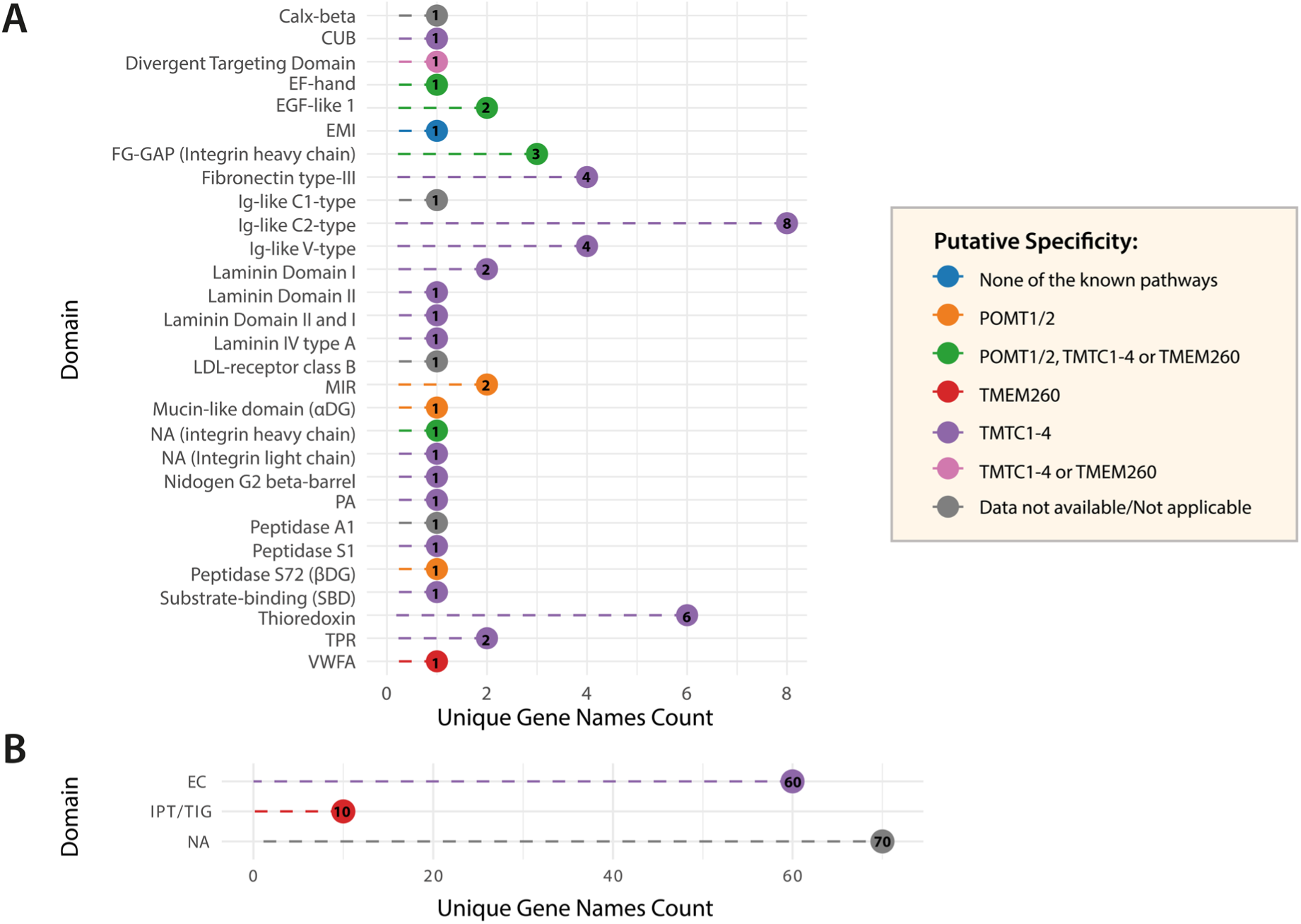
Protein domains targeted by O-Man glycosylation. **(A)** 30 protein domains are identified with O-Man glycosylation. O-Man deconstruction data suggest that TMTC1-4 have capacity to modify several domains and folds, while POMT1/2 and TMEM260 display activity limited to specific protein domains. **(B)** Number of canonical domains identified for TMTC1-4 and TMEM260 pathways. O-Man on unstructured regions on 70 distinct proteins is initiated by either of the three pathways (*cf.* Fig. 4A).

**Figure S7.**
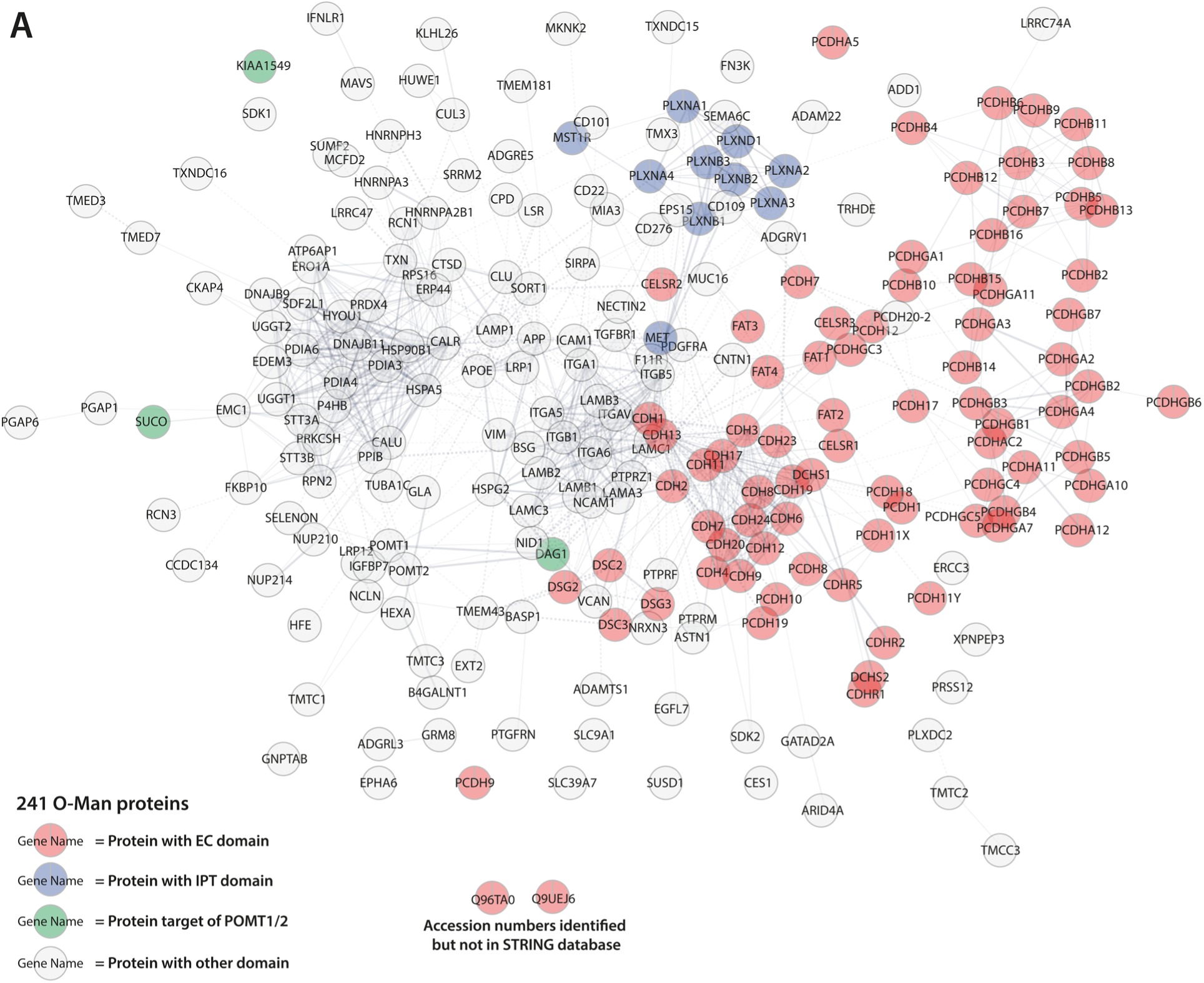
Overview of currently known O-Man proteins in humans. **(A)** Cumulative visualization of the 241 O-Man proteins identified in this study and in previous glycoproteomics studies (13,17–19,35) using STRING according to Methods section. In red are EC domains, in blue IPT domains, in green known POMT1/2 substrates and in gray are proteins with other domains. Few clusters are identified other than these two, and these involve laminins/integrins as well as other ER-resident proteins.

