## Supplementary Information for "Global view of domain-specific O-linked mannose glycosylation in glycoengineered cells"

### Supplementary material

Supplementary material containing datasets is uploaded as a separate Excel file.

Identification and quantification of relative abundances of glycopeptides enriched from crude membrane preparations, total cell lysates or from purified proteins. The experimental setup of paired (case and control) analyses of genetically engineered cell lines and their stable isotope coding is described under "Overview" sheet. Results from each analysis can be found in individual sheets (Dataset 1-36).

### Supplementary tables

#### Supplementary table 1 – gRNAs

| Gene name | gRNA Sequence (PAM) | gRNA plasmid ID | Addgene # | Reference |
|---|---|---|---|---|
| **POMT1** | GAGCTCCAACACTATCTGGT(**AGG**) | Gh23 | 106696 | Described previously in (13,59) |
| **POMT2** | CTTCGAGGCGGTCGGCTGGT(**GGG**) | Gh24 | 106697 | Described previously in (13,59) |
| **TMEM260** | TGGCTATCCTTTGTTCACGC(**TGG**) | Gh231 | N/A | Described previously in (18) |

#### Supplementary table 2 – Primers and oligoes

| Name | Sequence | Usage | Notes |
|---|---|---|---|
| **DAG1_cDNA_Fwd** | 5’ – ATGAGGATGTCT GTGGGCCTCTCGC TGCTG – 3’ | Amplification of DAG1 from HEK293 total cDNA | |
| **DAG1_cDNA_Rev** | 5’ - AGGTGGGACATA GGGAGGAGGTGA CCGG – 3’ | Amplification of DAG1 from HEK293 total cDNA | |
| **POMT1_IDAA_FwdExt** | 5′- AGCTGACCGGCA GCAAAATTGATC | Validation by IDAA of POMT1 KO cells | Described previously in (13) |

| | | | |
|---|---|---|---|
| | AACAGAGCAGCT CCCAT – 3’ | | |
| **POMT1_IDAA_Rev** | 5′- CATGACGGCGCT ATGTGAAA – 3’ | Validation by IDAA of POMT1 KO cells | Described previously in (13) |
| **POMT2_IDAA_FwdExt** | 5′- AGCTGACCGGCA GCAAAATTGCCT GGCAGAGTCCGA GCT -3’ | Validation by IDAA of POMT2 KO cells | Described previously in (13) |
| **POMT2_IDAA_Rev** | 5′- GACAGCAGCGTC ACCAAG – 3’ | Validation by IDAA of POMT2 KO cells | Described previously in (13) |
| **TMEM260_IDAA_FwdEext** | 5’- AGCTGACCGGCA GCAAAATTGCCA TGTGATAGACGCT GCCA-3’ | Validation by IDAA of TMEM260 KO cells | Described previously in (18) |
| **TMEM260_IDAA_Rev** | 5’- CATGTGTTAGGG AAACCAAGCAA - 3’ | Validation by IDAA of TMEM260 KO cells | Described previously in (18) |

### Supplementary table 3 – Cell lines generated and used

| Name of the cell line in this study | Parental cell line | Reference parental cell line | Modification to parental | INDEL generated and sequence of edited locus (or reference of original study) |
|---|---|---|---|---|
| **HEK293 or HEK293$^{WT}$** | N/A | N/A | N/A | N/A |
| **BG1** | N/A | N/A | N/A | N/A |
| **CaCo-2** | N/A | N/A | N/A | N/A |
| **HEPG2** | N/A | N/A | N/A | N/A |
| **SHSY-5Y** | N/A | N/A | N/A | N/A |
| **HEK293$^{SC}$** | HEK293$^{WT}$ | N/A | KO of *COSMC* and *POMGNT1* | Described in (13) |
| **HEK293$^{POMT1/2}$** | HEK293$^{KO:TMTC1-4}$ | (17) | KO of *TMEM260* | TMEM260: +1bp TGGCTATCCTTTGTT**T**CAC GC**TGG** |
| **HEK293$^{TMTC1-4}$** | HEK293$^{KO:POMT1, KO:POMT2}$ | (13) | KO of *TMEM260* | TMEM260: +1bp TGGCTATCCTTTGTT**T**CAC GC**TGG** |
| **HEK293$^{TMEM260}$** | HEK293$^{KO:TMTC1-4}$ | (17) | KO of *POMT1* and *POMT2* | POMT1: +2bp GAGCTCCAACACTATC**TG** TGGT**AGG** |

| | | | | |
|---|---|---|---|---|
| | | | | POMT2, allele 1: -1bp<br>CTTCGAGGCGGTCGGCTG<br>GTGGG<br>POMT2, allele 2: +81bp<br>CTTCGAGGCGGTCGGCT+<br>81bpGGTGGG |
| **HEK293$^{nO\text{-}Man}$** | HEK293$^{TMEM260}$ | This study | KO of *TMEM260* | TMEM260: +1bp<br>TGGCTATCCTTTGTTTCAC<br>GCTGG |
| **HEK293$^{WT+DAG1}$** | HEK293$^{WT}$ | N/A | ZFN KI of DAG1-3xFLAG | N/A |
| **HEK293$^{SC/SC+DAG1}$** | HEK293$^{SC/SC}$ | (13) | ZFN KI of DAG1-3xFLAG | N/A |
| **HEK293$^{nO\text{-}Man+DAG1}$** | HEK293$^{nO\text{-}Man}$ | This study | ZFN KI of DAG1-3xFLAG | N/A |

### Supplementary table 4 – Proteome Discoverer search parameters

| Dataset type (Main figure) | Precursor mass tolerance | Static Modifications / +Mass (residue) | Dynamic modifications | Quant |
|---|---|---|---|---|
| **Standard workflow (Fig. 1) – .raw from** (17) | 15 ppm | Carbamidomethyl / +57.021 Da (C)<br>Dimethyl:2H(4) / +32.056 Da (Any N-Terminus)<br>Dimethyl:2H(4) / +32.056 Da (K) | Oxidation / +15.995 Da (M)<br>Hex / +162.053 Da (S, T) | No |
| **Improved workflow (Fig. 1) –.raw from Figs. 3, 4, 6** | 15 ppm | Carbamidomethyl / +57.021 Da (C)<br>Diethyl:[1,2]13C2 / +60.076 Da (Any N-Terminus)<br>Diethyl:[1,2]13C2 / +60.076 Da (K) | Oxidation / +15.995 Da (M)<br>Hex / +162.053 Da (S, T) | No |
| **WT cells (Fig. 2, Fig. 6)** | 10 ppm | Carbamidomethyl / +57.021 Da (C)<br>Diethyl / +56.063 Da (Any N-Terminus)<br>Diethyl / +56.063 Da (K) | Oxidation / +15.995 Da (M)<br>Hex / +162.053 Da (S, T, W) | No |
| **Glycoengineered cells (Fig. 1 CMP and TCL, Fig. 3-4, Fig. 6)** | 10 ppm | Carbamidomethyl / +57.021 Da (C)<br>Diethyl:[1,2]13C2 / +60.076 Da (Any N-Terminus)<br>Diethyl:[1,2]13C2 / +60.076 Da (K)<br>Diethyl / +56.063 Da (Any N-Terminus)<br>Diethyl / +56.063 Da (K) | Oxidation / +15.995 Da (M)<br>Hex / +162.053 Da (S, T, W) | Yes |
| **Ratiochecks (Fig. S2)** | 15 ppm | Carbamidomethyl / +57.021 Da (C)<br>Diethyl:[1,2]13C2 / +60.076 Da (Any N-Terminus)<br>Diethyl:[1,2]13C2 / +60.076 Da (K)<br>Diethyl / +56.063 Da (Any N-Terminus)<br>Diethyl / +56.063 Da (K) | Oxidation / +15.995 Da (M) | Yes |
| **Soluble model proteins (Fig. 5)** | 10 ppm | Carbamidomethyl / +57.021 Da (C)<br>Diethyl:[1,2]13C2 / +60.076 Da (Any N-Terminus)<br>Diethyl:[1,2]13C2 / +60.076 Da (K)<br>Diethyl / +56.063 Da (Any N-Terminus)<br>Diethyl / +56.063 Da (K) | Oxidation / +15.995 Da (M)<br>Hex / +162.053 Da (S, T, W)<br>HexNAc / +203.079 Da (S, T) | Yes |

| | | | | |
|---|---|---|---|---|
| **Full-length DAG1 (Fig. S4)** | 10 ppm | Carbamidomethyl / +57.021 Da (C)<br>Diethyl:[1,2]13C2 / +60.076 Da (Any N-Terminus)<br>Diethyl:[1,2]13C2 / +60.076 Da (K)<br>Diethyl / +56.063 Da (Any N-Terminus)<br>Diethyl / +56.063 Da (K) | Oxidation / +15.995 Da (M)<br>Hex / +162.053 Da (S, T, W)<br>HexNAc / +203.079 Da (S, T) | No |
